# Bridge protein-mediated viral targeting of cells expressing endogenous μ-opioid G protein-coupled receptors in the mouse and monkey brain

**DOI:** 10.1101/2025.02.08.637063

**Authors:** Riki Kamaguchi, Satoko Amemori, Ken-ichi Amemori, Fumitaka Osakada

## Abstract

Targeting specific cell types is essential for understanding their functional roles in the brain. Although genetic approaches enable cell-type-specific targeting in animals, their application to higher mammalian species, such as nonhuman primates, remains challenging. Here, we developed a nontransgenic method using bridge proteins to direct viral vectors to cells endogenously expressing μ-opioid receptors (MORs), a G protein-coupled receptor. The bridge protein comprises the avian viral receptor TVB, the MOR ligand β-endorphin (βed), and an interdomain linker. EnvB-enveloped viruses bind to the TVB component, followed by the interaction of βed with MORs, triggering viral infection in MOR-expressing cells. We optimized the secretion signals, domain configurations, and interdomain linkers of the bridge proteins to maximize viral targeting efficiency and specificity. Alternative configurations incorporating different ligands and viral receptors also induced viral infection in MOR-expressing cells. The optimized βed-f2-TVB bridge protein with EnvB-pseudotyped lentiviruses induced infection in MOR-expressing cells in the striatum of mice and monkeys. An intersectional approach combining βed-f2-TVB with a neuron-specific promoter refined cell-type specificity. This study establishes the foundation for the rational bridge protein design and the feasibility of targeting G protein-coupled receptors beyond tyrosine kinase receptors, thereby expanding targetable cell types in the brain and throughout the body.

**Graphical abstract:** 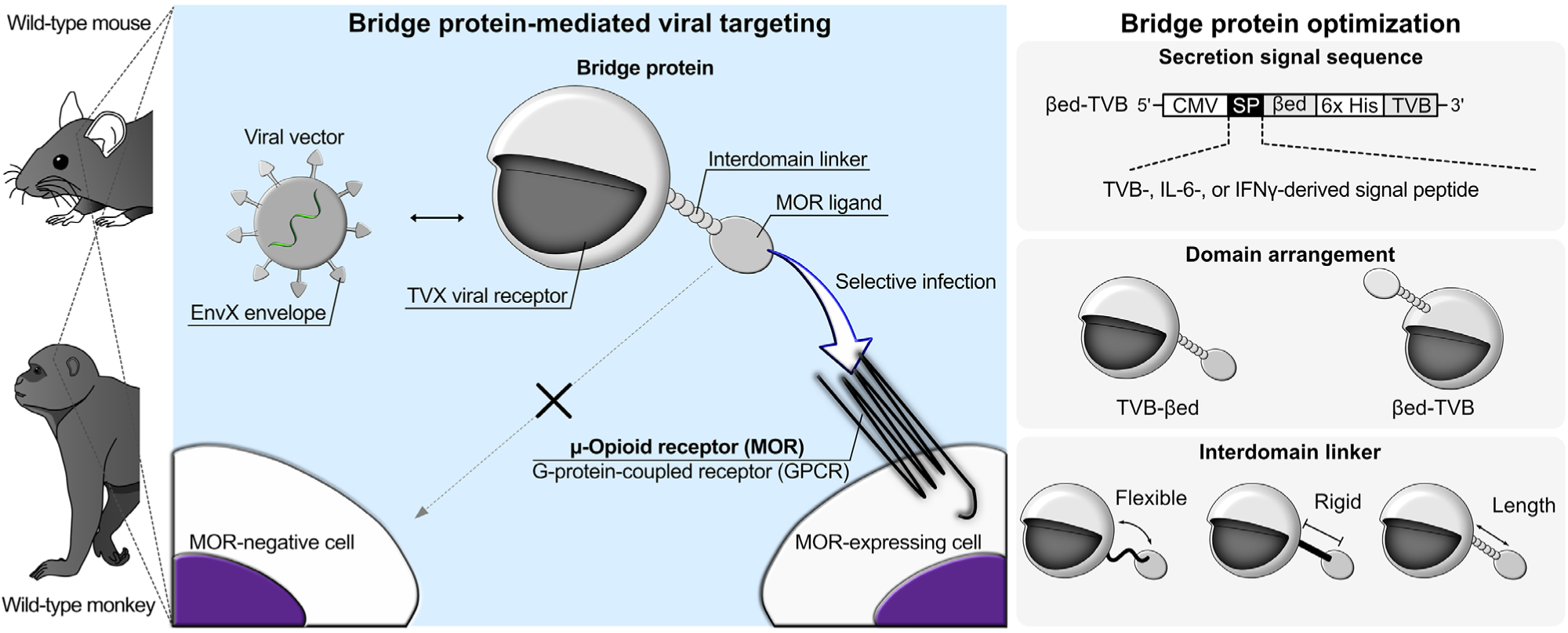

## Introduction

Neuroscience seeks to elucidate the mechanisms underlying brain function and to develop effective therapies for neurological and psychiatric disorders. The mammalian brain comprises billions of neurons and numerous non-neuronal cells, such as astrocytes, oligodendrocytes, and microglia, with extremely complex interactions through synaptic and non-synaptic connections among all these cells (Luo, 2021; Nagai et al., 2021). These cellular landscapes are further complicated by the recent findings that cell classes, such as neurons and astrocytes, are not homogeneous, but rather comprise diverse cell types, each distinguished by morphology, gene expression, function, and connectivity (Zeng, 2022). This cellular diversity underpins brain function and contributes to the pathogenesis of neurological and psychiatric disorders, with each cell type playing a distinct role in these processes (Adamczyk, 2023; Paolicelli et al., 2022; Sofroniew, 2020). Therefore, dissecting brain structure, function, and disease at the cell-type level is crucial for advancing our understanding of neural circuits and disease mechanisms.

In recent decades, notable advances have been made in experimental techniques for labeling and manipulating specific cell types in the mammalian brain (Luo et al., 2018). Cell-type classification is typically based on their electrophysiological properties, response characteristics, cell morphology, gene/protein expression patterns, and neuronal connectivity. In particular, molecular profiling through single-cell transcriptomes has revolutionized cell-type classification. This progress has led to the generation of transgenic animals expressing a particular gene of interest, including the recombinase Cre and Flp and the tetracycline transactivator tTA, in molecularly defined cell types, allowing studies linking the molecularly defined cell types to their functions in neural circuits and behavior (Atasoy et al., 2008; Branda and Dymecki, 2004; Huang and Zeng, 2013). Advances in genome-editing technologies, such as CRISPR-Cas9, have further facilitated the generation of transgenic animals (Horii et al., 2017; Ohtsuka et al., 2018). However, these approaches remain largely confined to a limited range of species, predominantly mice and rats. Extending these approaches to nonhuman primates (NHPs), carnivores (e.g., ferrets), or rodents (e.g., hamsters and guinea pigs) remains a notable challenge due to the difficulty of generating transgenic animals in these species (Kishi et al., 2014; Park and Silva, 2019; Shinmyo et al., 2017). To overcome this limitation, viral vector-based approaches have emerged as a powerful alternative for targeting and manipulating molecularly defined cell types in nontransgenic animals (Luo et al., 2018; Nectow and Nestler, 2020). Current viral approaches to target specific cell types include the use of promoters, enhancers, sesRNA, bridge proteins, and capsid engineering (Chen et al., 2022; Choi et al., 2010; Chuapoco et al., 2023; Dobson et al., 2022; Qian et al., 2022; Salimando et al., 2023). However, these viral targeting methods still face challenges in achieving high efficiency and specificity, especially in nontransgenic animals, such as NHPs.

One innovative approach involves using bridge proteins to direct enveloped viral vectors, such as lentiviruses and G-deleted rabies viruses, to particular cell types (Choi et al., 2010; Choi and Callaway, 2011). Bridge proteins are designed to link enveloped viruses to specific endogenous surface receptors on target cells, enabling the targeted viral infections that require no transgenic animals. Bridge proteins comprising neuregulin and TVB have been used to target ErbB4-expressing inhibitory neurons in the cerebral cortex (Choi et al., 2010; Choi and Callaway, 2011). In this system, EnvB-enveloped viruses bind to the TVB component of the bridge protein, followed by the interaction of the bridge protein ligand neuregulin-1 with endogenous ErbB4 receptors on the target cell surface. This interaction triggers selective viral infection of ErbB4-expressing inhibitory neurons. ErbB4 is a tyrosine kinase receptor that is internalized upon ligand binding. Bridge protein-mediated viral infection is thought to involve the internalization of surface receptors that bind to bridge proteins and interact with viral particles. However, it is unclear whether bridge proteins can target other classes of surface proteins, such as G protein-coupled receptors (GPCRs), ion channels, and transporters, which are widely expressed on cell membranes. GPCRs represent a promising target for the bridge protein system due to their well-characterized pharmacology and ability to undergo ligand-induced internalization (Mores et al., 2019). In addition, while the bridge protein system has great potential to target specific cell types, the design principles of bridge proteins, which are fundamental to guiding further development and improvement, remain lacking, limiting its broader applications.

In the present study, we investigated whether GPCRs on the cell membrane can serve as targets for bridge protein-mediated viral infection (Fig. 1A). We focused on μ-opioid receptors (MORs), a GPCR known to undergo ligand-induced internalization (Arttamangkul et al., 2019; Erbs et al., 2015; Gaudriault et al., 1997) and to show unique structural expression patterns in the striatum (Brimblecombe and Cragg, 2017; Johnston et al., 1990). To specifically target medium spiny neurons expressing GPCR MORs in the mammalian brain, we developed bridge proteins that comprise an MOR peptide ligand and a viral receptor and identified key drivers of the efficiency and specificity of bridge protein-mediated viral infection. We also explored different MOR ligands and various viral TVX receptors as a bridge protein configuration to expand the versatility of the bridge protein system. Our findings demonstrate that bridge protein-mediated viral targeting strategies can be successfully extended to GPCRs, beyond the previously established tyrosine kinase receptors. Integrating the bridge protein system based on receptors expressed on target cells with promoter/enhancer-based strategies in the viral genome enabled intersectional approaches to refine target cell specificity in nontransgenic animals, including NHPs. This study establishes a framework for the rational design of bridge proteins and expands the potential applications of bridge protein systems to various cell types in nontransgenic animals. By extending the bridge protein system to target a wider array of surface receptors, we paved the way for innovative strategies in targeted viral delivery and cellular manipulation.

**Fig. 1.**
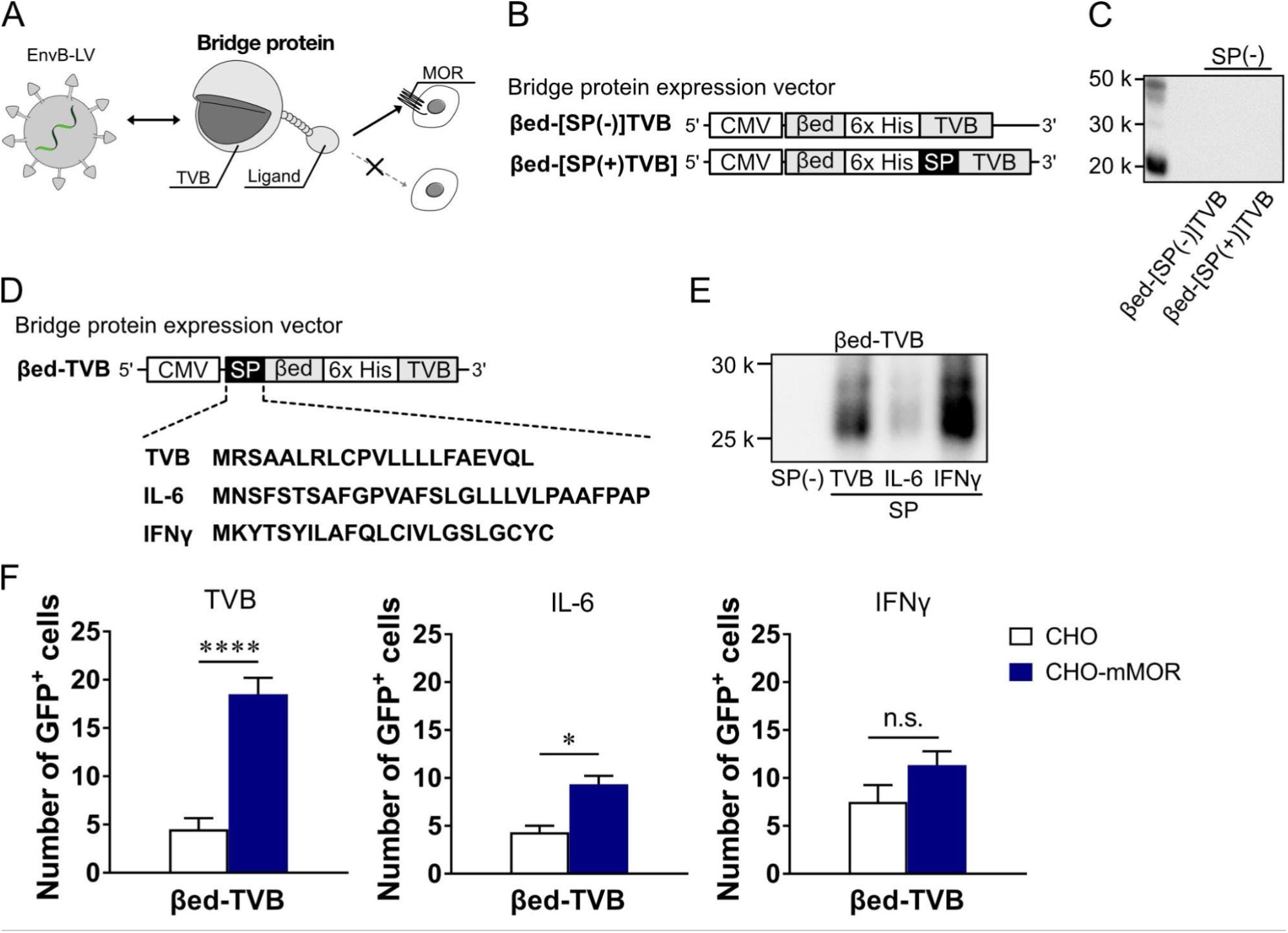
Optimization of secretion signal peptides in bridge proteins. (A) Schematic of the bridge protein system. (B) Bridge protein expression vectors with/without a secretion signal peptide sequence at the N terminus of the TVB. (C) Detection of secreted bridge proteins without the signal peptide by Western blotting with anti-Histidine tag antibody. (D) Schematic diagram of the bridge protein βed-TVB with the TVB-, IL-6-, or IFNγ-derived secretion signal peptide fused to the N terminus of βed. (E) Detection of the secreted bridge proteins with the TVB-, IL-6-, or IFNγ-derived signal peptide at the N terminus of βed. (F) Effects of bridge protein secretion signals on viral infection. EnvB-LV-CAG-EGFP was infected into CHO and CHO-mMOR cells in the presence of bridge proteins with TVB-(left), IL-6-(middle), or IFNγ-(right) derived secretion signal peptide. \**p* < 0.05, \*\*\*\**p* < 0.0001, vs. CHO (Unpaired *t*-test). Each column represents the mean ± SEM (n = 3−6).

## Materials and methods

### Animals

Mice were treated following with the Guidelines of Animal Experiments of Nagoya University. All animal experiments were approved by the Animal Care and Use Committee of Nagoya University. All efforts were made to reduce the number of animals used and minimize suffering and pain. Wild-type C57BL/6J mice were purchased from Nihon SLC (Shizuoka, Japan). The animals were housed in a temperature-controlled room (24°C ± 1°C) under a 12-h light/dark cycle with *ad libitum* access to food and water. Male and female mice older than 8 weeks were used in the experiments. A female *Macaca fuscata* monkey (8 years old, 5.6 kg) was also used in the experiments by following the Guidelines for Care and Use of Nonhuman Primates (3rd edition) with the approval of the Animal Experimental Committee of Kyoto University.

### Plasmid construction

DNA fragments were amplified by polymerase chain reaction (PCR) using PrimeSTAR Max DNA Polymerase (Takara) on a PCR thermal cycler (Takara). Plasmids digested by restriction enzymes and PCR fragments were assembled with NEBuilder HiFi DNA Assembly Master Mix (New England BioLabs). Chemically competent Stbl3 (Thermo Fisher Scientific) and XL10-Gold (Agilent Technology) *E. coli* cells were transformed with LV plasmids and rabies viral genome plasmids, respectively. Chemically competent DH5α *E. coli* cells (TOYOBO) were used for the bridge protein plasmids and other plasmids. Transformed *E. coli* cells were cultured in LB medium (Kanto Chemical) containing ampicillin (100 µg/ml).

### Production of HIV lentivirus

HIV-based lentivirus (LV) used for the *in vitro* infection assay was produced by HEK293T cells as described previously (Kodera et al., 2023). HEK293T cells were transfected with the LV genomic plasmid pBOB-CAG-EGFP, pBOB-hSyn-tTA-cHS4-tetO-EGFP, or pBOB-hSyn-tTA-cHS4-tetO-mScarlet3 and the packaging helper plasmids pMDL, pRSV, and pCI-EnvX using polyethylenimine (PEI) MAX (pH 7.0, Polysciences) at 37°C, 5% CO_2_ for 16 h and subjected to medium change with fresh one. Virus-containing supernatants were collected 40 h posttransfection.

For the animal experiments, the collected supernatants were centrifuged at 6,000 *g* for 24 h. After centrifugation, the viral pellets were resuspended in 4 ml of HBSS, loaded onto gradients (20% and 55%) of sucrose in phosphate-bufferd saline (PBS), and then ultracentrifuged at 200,000 *g* for 2 h (Beckman Coulter, Brea, CA, USA). After ultracentrifugation, the viral fraction was collected and concentrated using Amicon ultracentrifugal filters (Millipore). Virus aliquots were stored at −80°C until use. Infectious titers were determined using HEK-TVB cells. The titers of EnvB-LV-CAG-EGFP, EnvB-LV-hSyn-cHS4-tetO-EGFP, and EnvB-LV-hSyn-cHS4-tetO-mScarlet3 were 1.55-2.15 × 10^12^, 7.99 × 10^11^, and 7.59 × 10^11^ viral genome/ml, respectively.

### Production of oEnvB-pseudotyped RVΔG

Production of RVΔG-EGFP was described as previously (Osakada et al., 2011; Osakada and Callaway, 2013; Suzuki et al., 2020). Briefly, RVΔG-EGFP was recovered in B7GG cells by transfection with the viral genomic plasmid pSADΔG-EGFP and the plasmids encoding the viral proteins pcDNA-B19G, pcDNA-B19N, pcDNA-B19P, and pcDNA-B19L. During RVΔG-EGFP production, B7GG cells were maintained in DMEM supplemented with 1% FBS in a humidified atmosphere of 3% CO_2_ at 35°C. For pseudotyping RVΔG-EGFP with oEnvB, HEK-oEnvB cells were infected with unpseudotyped RVΔG-EGFP. The virus-containing medium was concentrated by ultracentrifugation at 70,000 *g* for 2 h (Beckman Coulter, Brea, CA, USA). The viral pellets were resuspended with HBSS, loaded onto gradients (20% and 55%) of sucrose in PBS, and ultracentrifuged at 200,000 *g* for 2 h, and a virus-containing layer was collected. The infectious titers of oEnvB-RVΔG-EGFP were determined using HEK293T and HEK-TVB cells. HEK293T cells were used to inspect contamination with unpseudotyped RVΔG-EGFP. The virus aliquots were stored at −80°C until use. The infectious titer of EnvB-RVΔG-EGFP used in the present study was 7.0 × 10^9^ infectious unit/ml.

### Bridge protein production

HEK293T cells were transfected with the bride protein-coding pcDNA-CMV-TVB-βed, pcDNA-CMV-βed-TVB, pcDNA-CMV-TVB-EM1, or pcDNA-CMV-EM1-TVB vectors and pAdVAntage (Promega) using PEI MAX (pH 7.0). After transfection, the cells were incubated at 33°C, 6% CO_2_ for 4 days, and their culture media were collected. The collected media were centrifuged at 2000 rpm for 3 min and used for Western blotting or the viral infection assay. For the animal experiments, the bridge proteins were purified with Ni Sepharose 6 Fast Flow resin (Cytiva). The corrected media were incubated with Ni resin equilibrated with Wash Buffer 1 (50 mM sodium phosphate buffer, 150 mM NaCl, 15 mM imidazole, pH 8.0) at 4°C overnight. After incubation, the resin was loaded on the column and washed with Wash Buffer 1, followed by Wash Buffer 2 (50 mM sodium phosphate buffer, 150 mM NaCl, 15 mM imidazole, 0.2% TritonX, pH 8.0), and then eluted from the column with Elution Buffer (50 mM sodium phosphate buffer, 500 mM NaCl, 100 mM imidazole). The fractions containing the purified bridge protein were determined by Western blotting. The solvent of the bridge protein was replaced with PBS and concentrated using Amicon Ultra (Millipore). The final concentration of the bridge protein solution was determined by a bicinchoninic acid (BCA) assay (Wako). The bridge protein solution was stored at −80°C.

### *In vitro* assay of bridge protein-mediated viral infection

For virus infection evaluation, the culture supernatants containing bridge protein and LV were collected after transfection and used. CHO-K1 cells stably expressing mouse MOR (CHO-mMOR cells) were generated using the PiggyBac system (Ding et al., 2005). The CHO-K1 cells were transfected with pPB-CAG-mMOR-IRES-BSD and pCAG-PBase and then selected with blasticidin (1000 μM). For all *in vitro* evaluations, HEK293T, CHO-K1, and CHO-mMOR cells were seeded in 48-well plates the day before the addition of the bridge protein/LV mixture solution. On the day after seeding, the collected culture supernatants containing bridge protein and LV (1/2-1/16 diluted) were mixed at a 1:1 volume ratio in low-binding tubes and incubated on ice for 1 h. After 1 h of incubation, the bridge protein/LV mixture solution was added to the 48-well plates and maintained at 35°C with 3% CO_2_ for 3 days. After 3 days of maintenance, the number of LV-derived GFP-positive cells in each well of the plate was counted for comparison.

### Immunoblotting

HEK293T cells were seeded in 6- or 10-cm dishes. After the transfection with a bridge protein-coding plasmid, the samples of the supernatant containing the bridge protein. Purified bridge proteins were mixed with the sample buffer (300 mM Tris-HCl (pH 6.8), 15% sodium dodecyl sulfate (SDS), and 60% glycerol) and boiled at 95°C for 5 min. Protein samples were subjected to sodium dodecyl sulfate–polyacrylamide gel electrophoresis (SDS-PAGE) on 15% polyacrylamide gels. Proteins were transferred from the SDS-PAGE gel to PVDF membranes (Immobilon, Merck Millipore). The membranes were blocked with Block-Ace (KAC) and labeled with horseradish peroxidase-conjugated anti-His IgG mouse polyclonal antibody (1:5000, Wako) overnight, 4°C. The membranes were treated with the chemiluminescent reagent ImmunoStar Zeta (Wako) and the immunopositive bands were detected with FUSION FX SPECTRA (Vilber).

### 3-(4,5-Dimethylthiazol-2-yl)-2,5-diphenyltetrazolium bromide (MTT) assay

Cell survival was evaluated using the MTT assay. CHO and CHO-mMOR cells were plated on 48-well plates at a density of 3.1 × 10^4^ cells/cm^2^. On the next day, the mixture of EnvB-LV-CMV-EGFP and βed-f2-TVB was applied to these cells. Three days after infection, cells were incubated in a medium containing 0.5 mg/ml MTT tetrazolium salt (Dojin) at 35°C in a 3% CO_2_ atmosphere for 4 h and then solubilized by addition of SDS for 18 h. The absorbance was measured at 570 nm using the Enspire plate reader (Perkin Elmer).

### Virus injection and perfusion

Adult C57BL/6J mice aged 8−12 weeks were anesthetized with a cocktail (10 ml/kg, intraperitoneally) containing medetomidine hydrochloride (75 µg/ml) and midazolam (400 µg/ml), followed by pentobarbital sodium salt (1 mg/ml). Three hours before the stereotaxic injection, EnvB-LV-CAG-EGFP was mixed with β-endorphin-TVB (βed-f2-TVB) at a 4:1 volume ratio between the virus and the purified bridge protein and incubated at room temperature (RT); 5-fold concentrations of the bridge proteins were used to achieve final concentrations of 1, 10, or 20 µM. After the reaction, 500 nl of this virus mixture was injected into the striatum (Anterior−Posterior: 0.64−0.92 mm, Medial−Lateral: ±1.19−1.61 mm, Dorsal-Ventral: 2.21-2.27 mm from the bregma) of the mice. Three weeks after injection, the animals were transcardially perfused with PBS and 4% paraformaldehyde in PBS. Brains were removed from the skull, postfixed with 4% paraformaldehyde in PBS for 1 h at RT, cryopreserved in 2% parafolmaldehyde/15% sucrose in PBS overnight, and then transferred to 30% sucrose in PBS, as described previously (Masaki et al., 2022; Okigawa et al., 2021). The brains were sectioned coronally at 40-µm thickness using a freezing microtome (REM-710; Yamato).

An adult Macaca fuscata monkey (8 years old, 5.6 kg, female) was anesthetized with an intramuscular application of dexmedetomidine (30 µg/kg), midazolam (150 µg/kg), and ketamine (5 mg/kg), followed by Isoflurane inhalation (1.0−2.5%). Three hours before the virus injection, EnvB-LV-CAG-EGFP or EnvB-LV-hSyn-tTA-cHS4-tetO-mScarlet3 was mixed with β-endorphin-f2-TVB at a 4:1 volume ratio with a final concentration of β-endorphin-f2-TVB of 20 µM, as in the experiments with mice. A Brainsight Vet Robot Neuronavigation System (Rogue Research, CA) with magnetic resonance and computed tomography images was used to determine the coordinates of the injection sites and to inject the bridge protein/EnvB-LV mixture. Viral mixtures at 5 µl were injected into the left side of the monkey’s striatum using a microinjection pump (UMP3; WPI, US) attached to the robot arm at the following coordinates: EnvB-LV-CAG-EGFP without βed-f2-TVB; AP: 19.707 mm, ML: 3.62 mm, DV: 23.715 mm, EnvB-LV-hSyn-tTA-cHS4-tetO-mScarlet3 without βed-f2-TVB; AP: 16.164 mm, ML: 4.688 mm, DV: 24.254 mm, EnvB-LV-CAG-EGFP with βed-f2-TVB; AP: 26.35 mm, ML: 4.368 mm, DV: 21.654 mm, EnvB-LV-hSyn-tTA-cHS4-tetO-mScarlet3 with βed-f2-TVB AP: 22.547 mm, ML: 2.506 mm, DV: 21.057 mm. Here, we determined 0 mm AP as the ear-bar 0, 0 mm ML as the middle point between the left and right edge of the ear bars, and 0 mm DV as the dura surface. The monkey was individually housed in a virus containment cage until the perfusion day according to the guideline for P2A level experiment of Kyoto University. Six weeks after the injection, the monkey was deeply anesthetized with the intravenous application of an overdose of thiopental sodium (25 mg/kg, *i.v.*), followed by heparin (10 IU/kg). Subsequently, the perfusion was made with 0.1 M PBS, followed by 4% PFA/PBS. The brain was removed from the skull, kept in 4% PFA/PBS for 24 h, and then replaced with 10, 20, and 30% of sucrose/PBS as it sank. The brain was blocked and covered with the OCT compound. The brain blocks were stored at −80°C. Coronal frozen sections were made at 15-µm thickness on a freezing microtome (REM-710; Yamato) and mounted on a glass slide.

### Immunohistochemistry with tyramide signal amplification

Tyramide signal amplification (TSA) for immunochemistry was conducted using a TSA kit (Invitrogen). Free-floating mouse brain sections were pre-treated with 3% H_2_O_2_ for 60 min to quench endogenous peroxidase activity. The sections were rinsed with PBS three times for 10 min and placed in blocking reagent for 2 h at RT (Yoshizawa et al., 2018). The sections were incubated with rabbit anti-MOR (1:10000, Immunostar) and chicken anti-GFP (1:1000, Abcam) antibodies at 4°C for 48 h. The sections were rinsed 3 times for 10 min with PBS containing 0.01% Triton X-100 (PBST) and incubated with horseradish peroxidase-labeled goat anti-rabbit IgG (1:1000, Invitrogen) at 4°C for 24 h. The sections were rinsed 6 times for 10 min in 0.01% PBST and reacted with biotinyl-tyramide (PerkinElmer) for 4 min at RT. The sections were rinsed 3 times for 10 min in 0.01% PBST and incubated with streptavidin-Alexa Fluor 594 (1:1000, Jackson ImmunoResearch, West Grove, PA, USA) and Alexa Fluor 488-conjugated anti-chicken IgY antibody (1:1000, Jackson ImmunoResearch) for 2 h at RT. The labeled cells were imaged with a confocal laser-scanning microscope with GaAsP detectors (LSM800, Zeiss) using a 10× (NA 0.45, Zeiss), 20× (NA 0.75, Zeiss), or 63× (NA 1.2, Zeiss) objective lens.

### RNAscope *in situ* hybridization

Monkey brain slides were stored at −80°C until they were ready for staining. Following the manufacturer’s protocol for the RNAscope Multiplex Fluorescent V2 assay (Advanced Cell Diagnostics), brain sections were stained with probes for EGFP (538851-C2), mScarlet3 (1299061-C2), and Oprm1 (518941-C3). Probes for EGFP and mScarlet3 were detected by Opal 570 (1:1500, Akoya Biosystems) and Oprm1 was detected by Opal 650 (1:1500). The slides were glass-covered using ProLong Gold (Thermo Fisher Scientific). Following the staining, the slides were imaged with a confocal laser-scanning microscope as described above for the mouse brain sections.

### Statistical analysis

Statistical analyses were performed using GraphPad Prism 7 (GraphPad, San Diego, CA, USA). Statistical significance was analyzed by unpaired *t*-test or one-way analysis of variance (ANOVA) followed by Tukey’s test or Dunnett’s test. The statistical analysis used is indicated in each figure legend. Probability values < 0.05 were considered statistically significant.

## Results

### Optimization of secretion signal sequences in bridge proteins

Bridge proteins require a peptide ligand that specifically binds to a receptor expressed on the target cells. To target MOR-expressing cells using bridge proteins, we used β-endorphin as the MOR peptide ligand. The design optimization of the bridge proteins involved systematically evaluating several critical parameters, including protein secretion signals, the order of fusion between the MOR ligand and the viral receptor TVB, and the type and length of the interdomain linkers connecting the ligand and TVB.

Protein secretion signals are essential for the efficient extracellular release, biosynthesis, folding kinetics, and stability of recombinant proteins (Brockmeier et al., 2006; Freudl, 2018). To produce bridge proteins comprising β-endorphin (βed) and TVB in HEK cells, we first evaluated the effect of the TVB secretion signal on bridge protein production. We generated two versions of β-endorphin and TVB bridge proteins with or without the TVB secretion signals at the N terminus of the TVB sequence: βed-SP(+)-TVB and βed-SP(−)-TVB (Fig. 1B). Both constructs included a 6× histidine tag for detecting and purifying the produced bridge proteins. Western blotting using an anti-His-tag antibody did not detect the presence of either bridge protein (βed-SP(+)-TVB or βed-SP(−)-TVB), regardless of the inclusion of the TVB secretion signal (Fig. 1C). These results suggest the need for further optimization of the secretion signal sequence in bridge protein production.

To improve bridge protein production and functionality, we systematically evaluated different secretion signals (Fig. 1D). In particular, we compared the secretion signal sequences derived from the native TVB peptide and two well-characterized secretion signals from interleukin-6 (IL6) and interferon gamma (IFNγ) (Han et al., 2017; Pan et al., 1987; Reinisová et al., 2008; Rinderknecht et al., 1984). In each construct, the secretion signal sequence was placed at the N terminus of the bridge protein to direct the bridge proteins to the secretory pathway, and a 6x histidine tag was included for detection and purification. In addition, TVB without the TVB secretion signal peptide was used to avoid the possibility of cleavage between βed and TVB. In the absence of the secretion signal, the bridge proteins were not detected in the culture media. In contrast, all three secretion signals from TVB, IL-6, and IFNγ facilitated bridge protein secretion into the culture media (Fig. 1E). The TVB- and IFNγ-derived signals were more efficient in secretion than the IL-6-derived signals. To evaluate the functionality of these secreted bridge proteins, we tested their ability to mediate viral infection in MOR-expressing cells (Fig. 1F). Each bridge protein was mixed with EnvB-pseudotyped lentiviral vectors carrying an EGFP reporter under the ubiquitous CAG promoter (EnvB-LV-CAG-EGFP), incubated on ice for 1 h, and then applied to CHO cells and CHO-mMOR cells, which stably express mouse MORs. GFP expression was observed in the CHO-mMOR cells, but little expression was observed in the CHO cells. Bridge proteins containing TVB- and IL6-derived signal peptides significantly increased the number of GFP-positive infected cells in CHO-mMOR cells compared with those with IFNγ-derived secretion signals. These results indicate that the choice of secretion signal sequence is crucial for designing efficient bridge proteins. The TVB secretion signals not only maximized protein production but also enhanced the ability of bridge proteins to mediate the selective virus infection of MOR-expressing cells. It is possible that the IL-6 signal peptide primarily affects protein production whereas the IFNγ signal negatively impacts protein activity, such as proper folding and stability. Therefore, we selected the TVB secretion signal for further TVB-based bridge protein designs in subsequent experiments.

### Type and length of interdomain linkers and ligand−receptor configuration in the bridge protein design

To optimize bridge protein functionality, we systematically investigated the effects of interdomain linker type and length, as well as the fusion order of the MOR ligand and the viral receptor TVB in the bridge proteins. Linker sequences are critical in determining the function of fusion proteins that connect two distinct protein domains, by influencing folding, stability, expression levels, bioactivity, and the interaction of the two fused domains (Chen et al., 2013). The linker flexibility and hydrophilicity can also significantly affect the functional integrity of the individual domains (Alfthan et al., 1995; Arai et al., 2001; Argos, 1990). In particular, the GS flexible linker exerts a positive effect on the folding and stability of various fusion proteins (Chen et al., 2013; Trinh et al., 2004).

We first compared the performance of the flexible and rigid linkers and the effect of domain arrangements in the bridge protein design. Four bridge protein configurations were generated, differing in the linker type and the orientation of βed and TVB (Fig. 2A). Bridge proteins with the βed ligand at either the N- or C terminus with a flexible or rigid linker were tested for viral infection. The secretion of each bridge protein into the culture media was detected by Western blotting (Fig. 2B). We then determined whether these bridge proteins could mediate the infection of MOR-expressing cells by EnvB-LV-CAG-EGFP *in vitro*. EnvB-LV-CAG-EGFP combined with each bridge protein induced GFP expression in CHO-mMOR cells, with little expression in CHO cells, suggesting MOR-specific viral infection (Fig. 2C). We found no significant difference in viral infection efficiency between flexible and rigid linkers. However, among the four designs, the configuration with βed at the N terminus and a flexible linker (βed-flexible-TVB) showed the highest infection efficiency in MOR-expressing cells. For both linkers, βed positioned at the N terminus of the bridge proteins consistently resulted in higher infection efficiency in MOR-expressing cells than βed at the C terminus (Fig. 2D, E). This result corroborates previous findings that peptide ligands have MOR-binding motifs at their N terminus (Gaudriault et al., 1997). Based on these results and previous reports demonstrating that flexible linkers generally enhance hydrophilicity and purification efficiency as well as minimize interference between functional domains, we selected flexible linkers for all subsequent bridge proteins. Together, we used the MOR ligand βed at the N terminus and the viral receptor TVB at the C terminus connected by a flexible linker for the bridge proteins, hereafter.

**Fig. 2.**
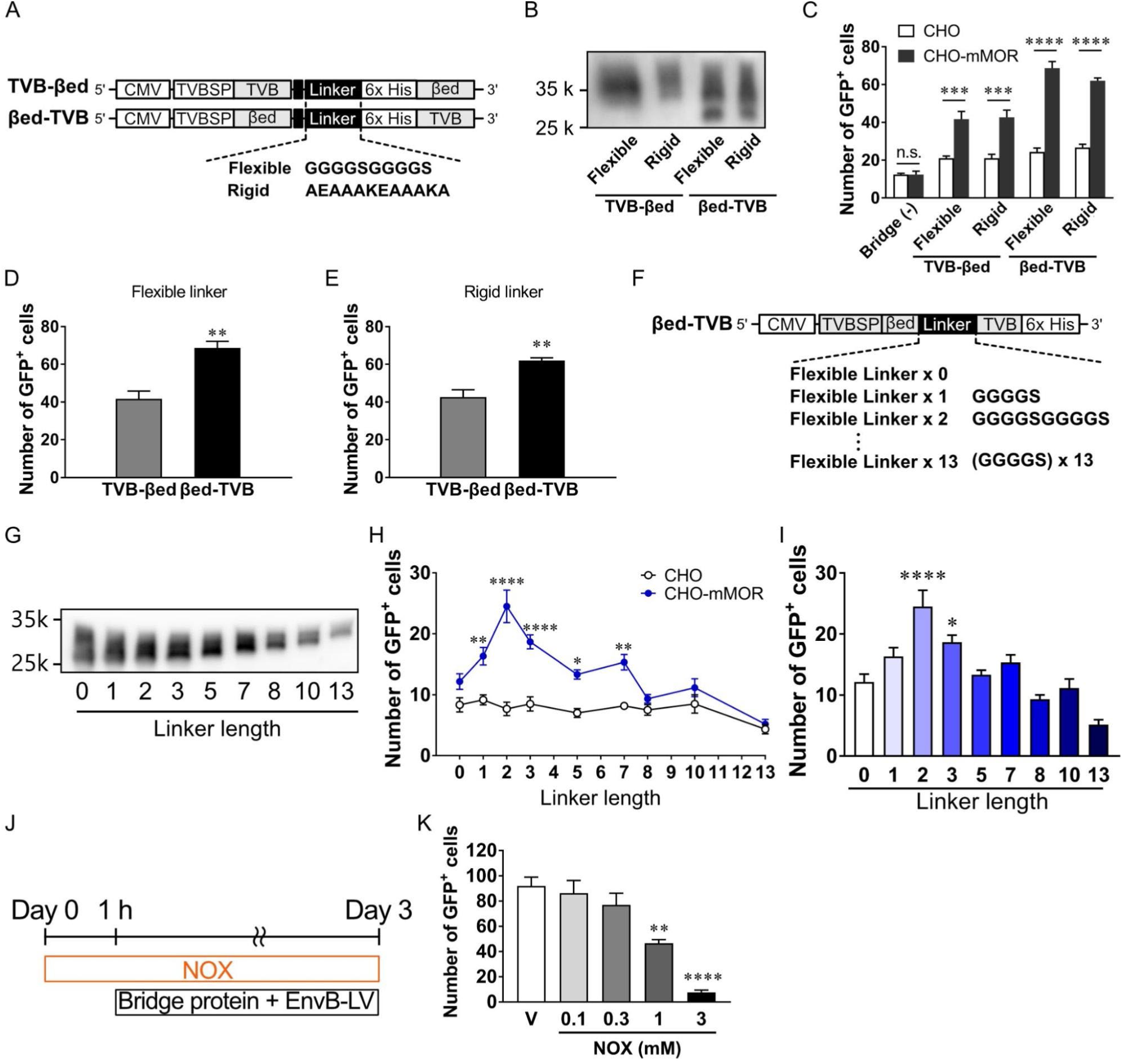
Type and length of interdomain linkers and domain order in bridge protein configurations. (A) Schematic diagram of the bridge protein expression vectors. Two types of flexible and rigid linkers were inserted between βed and TVB. A proline-rich linker was placed in front of a flexible or rigid linker and described as a black square (Snitkovsky et al., 2001). (B) Detection of secreted TVB-βed and βed-TVB with a flexible or rigid linker by Western blotting with anti-Histidine tag antibody. (C) Bridge protein-mediated viral infection efficiency with four types of βed-based bridge proteins. EnvB-LV-CAG-EGFP was infected into CHO and CHO-mMOR cells in the presence or absence of bridge protein. \*\*\**p*<0.001, \*\*\*\**p*<0.0001, vs. CHO (one-way ANOVA with Tukey’s test). Each column represents the mean ± SEM (n = 3). (D) Effect of ligand-receptor configuration of flexible-linker bridge proteins on viral infection. TVB-βed and βed-TVB using a flexible linker were used for the infection assay. \*\**p*<0.01, vs. TVB-βed (Unpaired *t*-test, n =3). (E) Effect of ligand−receptor configuration of rigid-linker bridge proteins on viral infection. TVB-βed and βed-TVB using a rigid liner were used for the infection assay. \*\**p*<0.01, vs. TVB-βed (Unpaired *t*-test, n = 3). (F) Different lengths of flexible interdomain linkers of the bridge protein. (G) Expression of βed-TVB with different flexible-linker lengths was detected by Western blotting with anti-Histidine tag antibody. (H) Comparison of viral infection efficiency using a bridge protein with different linker lengths. \**p*<0.05, \*\**p* < 0.01, \*\*\*\**p* < 0.0001, vs. CHO (one-way ANOVA with Tukey’s test, n = 6). (I) Linker length comparison with linker length 0 in CHO-mMOR. \**p*<0.05, \*\*\*\**p* < 0.0001, vs. Linker length 0 (one-way ANOVA with Dunnett’s test, n = 6). (J) Schematic of the pharmacological inhibition assay. The MOR antagonist Naloxone (NOX) at 0.1−3 mM was applied 1 h before and during the application of βed-f2-TVB and EnvB-LV-CAG-EGFP. (K) Concentration-dependent competitive inhibition of βed-f2-TVB-mediated viral infection by the MOR antagonist naloxone (NOX). \*\**p*<0.01, \*\*\*\**p* < 0.0001, vs. vehicle (one-way ANOVA with Dunnett’s test, n = 3).

Moreover, the interdomain linker length of a bifunctional protein alters the spatial arrangement of its functional domains, thereby affecting fusion protein stability and activity (Arai et al., 2001; Bai and Shen, 2006). To investigate the effect of interdomain linker length on bridge protein performance, we generated bridge proteins with varying lengths of interdomain linkers, defined by the number of repeating GGGGS units (Fig. 2F). Western blot analysis revealed bridge proteins with different linger lengths were detected at different molecular weight positions corresponding to the number of linker repeats (Fig. 2G). We then evaluated viral infection in CHO and CHO-mMOR cells using bridge proteins with different linker lengths. Viral infection was effective when the linker length was shorter than seven repeats (Fig. 2H). Notably, among the different linker lengths tested, the configuration with two flexible-linker repeats (βed-f2-TVB) induced the largest number of GFP-expressing cells, indicating that the two repeats provided the most efficient viral infection (Fig. 2I). These results indicate that the interdomain linker length plays a critical role in the bridge protein-mediated viral targeting of MOR-expressing cells.

To determine whether viral infection with βed-f2-TVB was mediated by MOR, we assessed the effect of the MOR antagonist naloxone (NOX) on the viral infection (Fig. 2J). The number of GFP-expressing cells infected with EnvB-LV-CAG-GFP, in the presence of βed-f2-TVB, significantly decreased in a concentration-dependent manner under NOX treatment at 1 and 3 mM. (Fig. 2K). This competitive blockade of βed-f2-TVB-mediated viral infection by NOX suggests that the viral infection was mediated by the bridge protein−MOR interaction on the target cells.

Thus, the configuration of the MOR ligand βed at the N terminus, the viral receptor TVB at the C terminus, and a two-repeat flexible linker (βed-f2-TVB) was identified as the optimal design for the bridge protein-mediated targeting of MOR-expressing cells. These results highlight the critical importance of ligand-receptor orientation and interdomain linker properties in optimizing the efficiency and specificity of bridge protein-mediated viral transduction.

### Combination of alternative ligands and viral receptors for bridge proteins

We explored whether alternative MOR ligands and viral receptors could be used in the bridge protein design to target MOR-expressing cells. Endomorphin-1 (EM1) (McConalogue et al., 1999), a peptide ligand with higher selectivity for MOR compared to βed, was selected as a candidate MOR ligand. We generated bridge proteins with four different configurations of the EM1 ligands positioned at either the N- or C terminus and linked by either a flexible or rigid linker (Fig. 3A). We then assessed the infectivity of these four bridge proteins in both the CHO and CHO-mMOR cells. EnvB-LV-CAG-EGFP in the presence of bridge protein induced GFP expression in CHO-mMOR cells (Fig. 3B). Regarding ligand orientation, placing the EM1 ligand at the N terminus resulted in a higher infection efficiency than that at the C terminus for both flexible and rigid linker types (Fig. 3C, D). In addition, when comparing the linker type, EM1-flexible-TVB showed greater efficiency in infecting MOR-expressing cells than EM1-rigid-TVB (Fig. 3E). Among the four configurations, EM1 positioned at the N terminus with a flexible linker (EM1-flexible-TVB) showed the highest infection efficiency in CHO-mMOR (Fig. 3B). Consistent with βed-f2-TVB, the optimal design of EM1-based bridge proteins featured the MOR ligand at the N terminus, TVB at the C terminus, and a flexible linker.

**Fig. 3.**
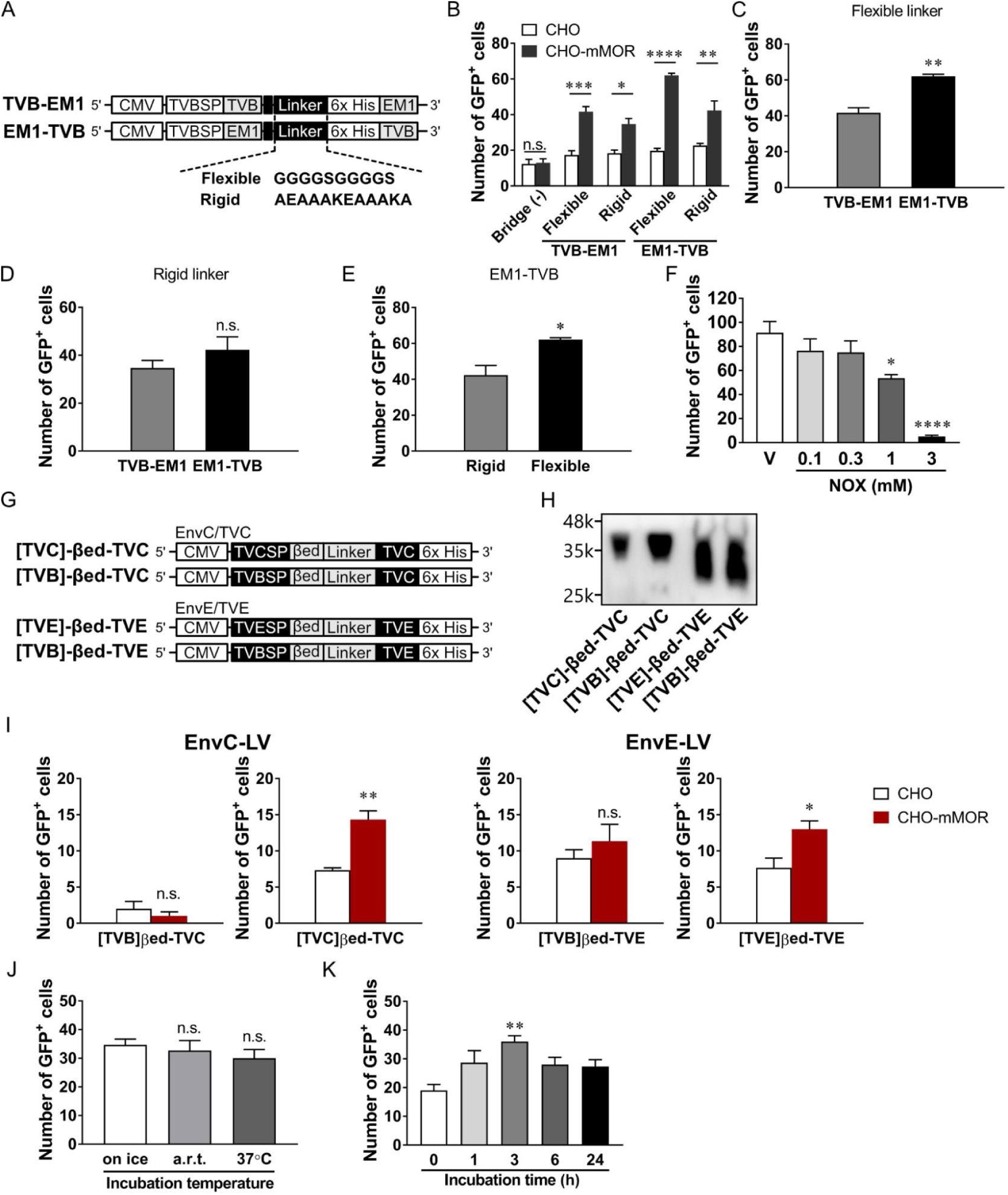
Different ligands and viral receptors for bridge protein design. (A) Expression vectors of the EM1 ligand-based bridge proteins. The TVB and EM1 were connected by a flexible or rigid linker of a different order. (B) Viral infection efficiencies with four types of EM1-based bridge proteins. \**p*<0.05, \*\**p*<0.01, \*\*\**p*<0.001, \*\*\*\**p*<0.0001, vs. CHO (one-way ANOVA with Tukey’s test). Each column represents the mean ± SEM (n = 3). (C) Effect of ligand−receptor configuration of flexible linker bridge proteins on viral infection. TVB-EM1 and EM1-TVB using a rigid linker were used for the infection assay. \*\**p*<0.01, vs. TVB-EM1 (Unpaired *t*-test, n = 3). (D) Effect of ligand-receptor configuration of rigid linker bridge proteins on viral infection. TVB-EM1 and EM1-TVB using a rigid liner were used for the infection assay. (E) Effect of EM1-TVB linkers on viral infection. EM1-TVB connected with a rigid or flexible linker was used for the infection assay.\**p*<0.05, vs. Rigid (Unpaired *t*-test, n = 3). (F) Concentration-dependent competitive inhibition of EM1-f2-TVB-mediated viral infection by the MOR antagonist. Naloxone (NOX) at 0.1−3 mM was applied 1 h before and during the application of the mixture of EM1-f2-TVB and EnvB-LV-CAG-EGFP. \**p*<0.05, \*\*\*\**p*<0.0001, vs. vehicle (one-way ANOVA with Dunnett’s test, n =3). (G) Design of the TVC- and TVE-based bridge proteins. Different subgroup receptors (TVC and TVE) to the corresponding avian virus envelopes (EnvC and EnvE) were connected to the MOR ligand βed with two repeats of a flexible linker. Secretion signal peptides from TVB (control) and endogenous TVC or TVE were used to produce the bridge proteins βed-TVC and βed-TVE. (H) Production of βed-TVC and βed-TVE using different secretion signals. βed-TVX bridge proteins with the corresponding endogenous secretion signal peptide or TVB-derived signal peptide (control) were detected by Western blotting with anti-Histidine tag antibody. (I) Viral infection via bridge proteins using the EnvC/TVC (left) and EnvE/TVE (right) systems. \**p*<0.05, \*\**p*<0.01, vs. CHO (Unpaired *t*-test, n = 3). (J) Effect of the incubation temperature of the bridge protein and virus mixture solution on infection efficiency. βed-TVB and EnvB-LV-CAG-EGFP were reacted on ice, at room temperature, or at 37°C before infection (one-way ANOVA with Dunnett’s test, n = 6). (K) Effect of the incubation time of βed-TVB and EnvB-LV-CAG-EGFP mixture solution. A mixture of βed-f2-TVB and EnvB-LV-CAG-EGFP was incubated on ice for 0, 1, 3, 6, and 24 hours before application to CHO-mMOR cells. \*\**p*<0.01, vs. 0 h (one-way ANOVA with Dunnett’s test, n = 6).

To evaluate the MOR specificity of EM1-f2-TVB, we performed a competitive inhibition assay using the MOR antagonist NOX (Fig. 2J). EM1-f2-TVB with EnvB-LV-CAG-EGFP increased the number of GFP-expressing cells in the CHO-mMOR. The number of GFP-expressing cells was reduced when NOX was applied 1 h before and during virus infection (Fig. 3F). The suppressive effect of NOX was concentration-dependent and reached statistical significance at concentrations of 1 and 3 mM. These results indicate that EM1-f2-TVB mediates the viral infection of MOR-expressing cells via EM1−MOR binding on the target cells. Notably, the most effective configuration, with a MOR ligand at the N terminus and TVB at the C terminus connected by a flexible linker, emerged as a common structural feature in bridge proteins for efficient viral targeting of MOR-expressing cells.

We next tested the feasibility of using an alternative envelope-receptor system of avian virus subgroups in the bridge protein design. We generated bridge proteins incorporating the EnvC/TVC and EnvE/EnvB-DR46 (TVE) system-derived receptors (Barnard et al., 2006; Barnard and Young, 2003; Matsuyama et al., 2015; Suzuki et al., 2020). Based on our results on TVB-based bridge proteins (Fig. 2F), the MOR ligand βed was positioned at the N terminus of the bridge protein, and a flexible linker with two repeats was used. For the TVC- and TVE-based bridge proteins, the extracellular domains of TVC and TVE were used for their receptor-binding domains (Fig. 3G), and their corresponding secretion signals were used. The TVB-derived secretion signal was utilized as a control. After the construction of plasmids encoding each bridge protein, the plasmids were transfected into HEK293T, and the secreted proteins containing the supernatant were collected. Western blot analysis revealed successful secretion of each bridge protein, regardless of the secretion signals (Fig. 3H). We evaluated the infection of βed-f2-TVC with EnvC-pseudotyped LV vectors in CHO and CHO-mMOR cells (Fig. 3I). The mixture of βed-f2-TVC with EnvC-LV-CAG-EGFP induced GFP expression in CHO-mMOR cells, with little GFP expression in CHO cells. The βed-f2-TVC bridge protein with TVB-derived secretion signals failed to induce MOR-specific infection, despite extracellular secretion into the culture media. In contrast, βed-f2-TVC with the TVC-derived secretion signal peptide provided efficient viral infection in MOR-expressing cells (Fig. 3I). We also examined the infection of MOR-expressing cells with βed-f2-TVE and EnvE-pseudotyped LV vectors. Viral infection assays showed that βed-f2-TVE with the EnvE-LV-CAG-EGFP induced GFP expression in CHO-mMOR cells, with some expression in CHO cells (Fig. 3I). These results indicate that βed-f2-TVE mediates viral infection in MOR-expressing cells. Overall, we conclude that alternative avian virus subgroups, such as the EnvC/TVC and EnvE/TVE systems, are applicable for the bridge protein-mediated targeting of MOR-expressing cells.

Beyond optimizing the design of bridge proteins to enhance viral infection efficiency, we investigated the effects of incubation time and temperature on bridge protein−virus interactions (Boerger et al., 1999; Mothes et al., 2000; Smith et al., 2004). Mixtures of βed-f2-TVB and EnvB-LV-CAG-EGFP were incubated for 1 h under three different temperature conditions, on ice, RT, and 37°C before being applied to CHO-mMOR cells. GFP expression was observed in the CHO-mMOR cells under all temperature conditions, but no statistical differences were observed in the number of virus-derived GFP-expressing cells between temperatures (Fig. 3J). We next examined whether varying the incubation time after mixing the bridge protein and viruses affected viral infection efficiency. Mixtures of βed-f2-TVB and EnvB-LV-CAG-EGFP were incubated on ice for various durations, 0, 1, 3, 6, and 24 h before being applied to CHO-mMOR cells. GFP expression was detected at all time points. The 3 h incubation provided the highest efficiency in the number of GFP-expressing cells with statistical significance compared with 0 h incubation (Fig. 3K). These results indicate that the incubation time for bridge protein−virus interaction is a critical factor in the performance of bridge protein systems. Based on these results, we adopted a 3 h incubation of bridge proteins and viral vectors at RT before infection as the standard protocol for subsequent experiments.

### Validation of bridge protein to MOR-expressing cells in nontransgenic mice *in vivo*

The striatum mainly comprises medium spiny neurons, which are inhibitory GABAergic neurons that constitute approximately 90% of its cellular population. The striatum is divided into two functional compartments: the striosome and the matrix, each characterized by distinct gene expression profiles (Graybiel and Matsushima, 2023). Despite the differences in their expression patterns between species, MORs are abundantly expressed in the medium spiny neurons of the striosomes but rarely in the matrix of the mouse striatum (Delfs et al., 1994; Mansour et al., 1994; Minami and Satoh, 1995). Based on this expression pattern, we sought to target MOR-expressing cells in the striatum using our developed bridge protein, βed-f2-TVB. To evaluate the *in vivo* specificity of the βed-f2-TVB bridge protein for MOR-expressing cells, we injected a mixture of βed-f2-TVB and EnvB-LV-CAG-EGFP into the striatum of wild-type mice (Fig. 4A, B). For *in vivo* validation, we used a lentiviral vector expressing transgenes under the ubiquitous CAG promoter. In addition, the efficiency in virus infection of target cells was concentration-dependent on bridge proteins *in vitro (Balliet et al., 1999)*. We therefore examined concentrations of bridge proteins necessary to achieve efficient infection and specificity to MOR-expressing cells *in vivo*. The mixture of βed-f2-TVB at concentrations of 1, 10, and 20 µM and EnvB-LV-CAG-EGFP were incubated for 3 h at RT and then injected into the striatum of wild-type mice. Three weeks after injection, GFP expression derived from EnvB-LV-CAG-EGFP was observed in the striatum at all tested bridge protein concentrations. In the presence of βed-f2-TVB, GFP-expressing cells were evident around the injection site of the striatum, whereas little EGFP expression was detected in the absence of the bridge protein (Fig. 4C). Notably, the number of GFP-positive cells increased in a concentration-dependent manner with βed-f2-TVB (Fig. 4D). To determine whether bridge protein-mediated viral infection was specific to MOR-expressing cells, we performed immunohistochemistry for GFP and MOR. The percentage of cells co-expressing EGFP and MOR increased with higher concentrations of βed-f2-TVB (Fig. 4E) and reached 44.8% at 20 μM. These results demonstrate that the βed-f2-TVB bridge protein directs EnvB-pseudotyped LV infection to MOR-expressing cells *in vivo*, albeit with moderate specificity.

**Fig. 4.**
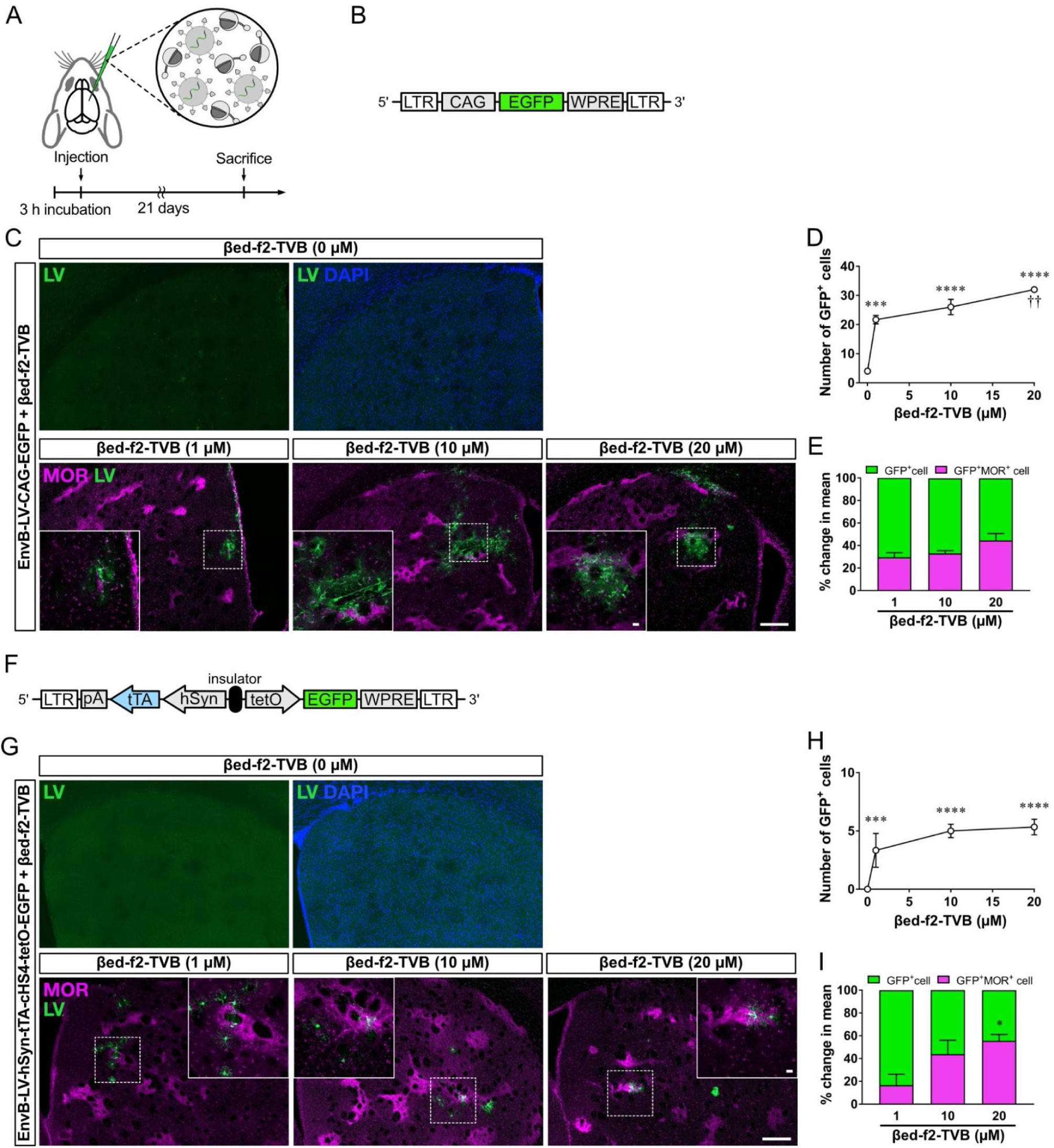
Bridge protein-mediated viral targeting of MOR-expressing cells in wild-type mice. (A) Experimental procedures for *in vivo* evaluation. A mixture of βed-f2-TVB and EnvB-LV-CAG-EGFP was injected into the striatum of wild-type C57BL6/J mice after incubation for 3 h at room temperature. The mice were sacrificed 3 weeks postinjection and processed for histological analysis. (B) Construct design for EnvB-pseudotyped lentiviral vectors encoding EGFP regulated by the CAG promoter. (C) Infection of EnvB-LV-CAG-EGFP with different concentrations of βed-f2-TVB in the mouse striatum. The white square indicates a representative high-magnification view of the white dotted line area. Scale bar: 200 and 20 µm (large and small panels). (D) Concentration-dependency of βed-f2-TVB in EnvB-LV-CAG-EGFP infection. \*\*\**P* < 0.001, \*\*\*\**p* < 0.0001, vs. Bride protein at 0 µM. ††*p* < 0.01, vs. Bridge protein at 1 µM (One-way ANOVA with Dunnett’s test, n = 3). (E) MOR-positive ratio in βed-f2-TVB-mediated EnvB-LV-CAG-EGFP-infected GFP-positive cells. (F) Construct design for EnvB-LV-hSyn-tTA-cHS4-tetO-EGFP. EGFP expression is self-amplified in neurons by the tetO promoter driven by tTA under the control of the neuron-specific human synapsin promoter. (G) EnvB-LV-hSyn-tTA-cHS4-tetO-EGFP with different concentrations of βed-f2-TVB. The white square indicates a representative high-magnification view of the white dotted line area. Scale bar: 200 and 20 µm (large and small panels). (H) Concentration-dependency of βed-f2-TVB in EnvB-LV-hSyn-tTA-cHS4-tetO-EGFP infection. \*\*\**p* < 0.001, \*\*\*\**p* < 0.0001, vs. Bride protein at 0 µM (one-way ANOVA with Dunnett’s test, n = 3). (I) MOR-positive ratio in βed-f2-TVB-mediated, EnvB-LV-hSyn-tTA-cHS4-tetO-EGFP-infected GFP-positive cells. \**p* < 0.05, vs. Bridge protein at 1 µM (one-way ANOVA with Dunnett’s test, n = 3).

Next, we sought to enhance the specificity of viral targeting to MOR-expressing neurons. MOR expression is not exclusive to neurons and is also observed in microglia (Maduna et al., 2018). In addition, microglia may phagocytose viral particles, leading to nonspecific GFP expression. The CAG promoter used in our LV vector, EnvB-LV-CAG-EGFP, drives transgene expression ubiquitously across various cell types. To overcome these limitations and improve the specificity of bridge proteins, we introduced a neuron-specific promoter in the LV vector. We designed an LV vector carrying a neuron-specific promoter coupled with a self-amplification system under the tetO promoter (EnvB-LV-hSyn-tTA-cHS4-tetO-EGFP), to achieve an intersectional approach based on MOR expression and neuronal specificity (Fig. 4F). The self-amplification vector expresses transgenes under the tetO promoter, which is activated by tTA expressed under the control of the neuron-specific human synapsin promoter. The βed-f2-TVB bridge protein at 1, 10, and 20 µM was mixed with EnvB-LV-hSyn-tTA-cHS4-tetO-EGFP, incubated for 3 h at RT, and then injected into the striatum of wild-type mice. GFP expressions were not detected in the absence of βed-f2-TVB. In contrast, co-injection of EnvB-LV-hSyn-tTA-cHS4-teOt-EGFP and βed-f2-TVB induced GFP expression in the striatum (Fig. 4H). Immunohistochemistry revealed that some GFP-expressing cells co-expressed MOR. The percentage of GFP and MOR double-positive cells increased at higher concentrations of the bridge protein. At 20 μM βed-f2-TVB, 56.3% of the GFP-expressing cells were positive for MOR (Fig. 4I). This MOR specificity by EnvB-LV-hSyn-tTA-cHS4-tetO-EGFP was higher than that by the CAG promoter-based system of EnvB-LV-CAG-EGFP (44.8%). Thus, this intersectional approach combining neuron-specific promoters with bridge proteins can improve the specificity of viral targeting to MOR-expressing neurons (Hioki et al., 2009; Watakabe et al., 2012).

### Transduction to MOR-expressing cells using G-deleted rabies virus vectors

In the next set of experiments, we tested the versatility of MOR-targeted bridge protein-mediated viral infection to another enveloped virus, such as G-deleted rabies virus vectors (RVΔG). We used the optimized βed-f2-TVB bridge protein, developed for LV (Fig. 2) for EnvB-pseudptyped (EnvB-RVΔG). We mixed EnvB-RVΔG-EGFP with βed-F2-TVB on ice for 1 h and then applied the mixture solution to CHO and CHO-mMOR cells. EnvB-RVΔG-EGFP-derived GFP-positive cells were detected in CHO-mMOR cells, but not CHO cells, suggesting that the bridge protein βed-F2-TVB mediated efficient and specific EnvB-RVΔG infection in MOR-expressing cells (Fig. 5A). Next, we investigated whether EnvB-RVΔG-EGFP mixed with the bridge protein βed-F2-TVB could infect MOR-expressing cells in the mouse brain. A mixture of EnvB-RVΔG-EGFP and βed-F2-TVB was incubated for 3 h at RT and injected into the striatum of wild-type mice (Fig. 5B). One week after injection, EnvB-RVΔG-EGFP-infected, GFP-expressing cells were detected in the striatum at 20 µM βed-f2-TVB, whereas little to no GFP expression was observed under βed-f2-TVB absence (Fig. 5C, D). We also determined the specificity of the bridge protein-mediated EnvB-RVΔG-EGFP infection in the MOR-expressing cells by immunohistochemistry. Among GFP-positive cells, 53.8% co-expressed MORs (Fig. 5E). These results indicate that the MOR-targeting bridge protein applies to EnvB-pseudotyped viral vectors, which corroborates a previous study using a TVB-NRG1 bridge protein (Choi et al., 2010).

**Fig. 5.**
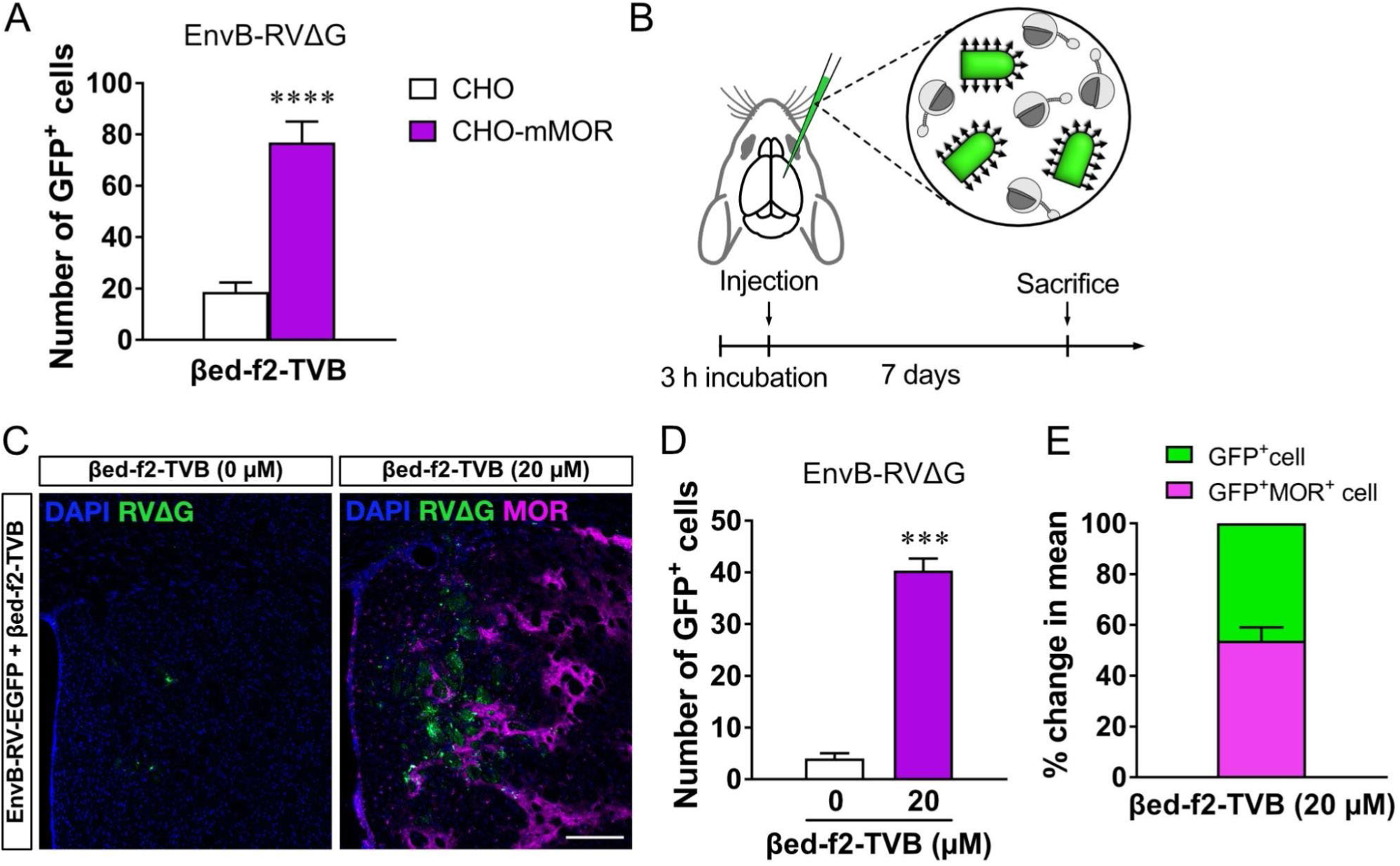
Bridge protein-mediated infection of G-deleted rabies virus vectors in MOR-expressing cells. (A) Bridge protein-mediated targeting of MOR-expressing cells with G-deleted rabies virus vectors. A mixture of EnvB-RVΔG-EGFP and the bridge protein βed-f2-TVB was applied to CHO and CHO-mMOR cells. \*\*\*\**p* < 0.0001, vs. CHO (Unpaired *t*-test, n = 3). (B) Experimental procedures for *in vivo* evaluation. A mixture of βed-f2-TVB and EnvB-RVΔG-EGFP was injected into the striatum of wild-type C57BL6/J mice after incubation for 3 h at RT. (C) *In vivo* infection of EnvB-RVΔG-EGFP with βed-f2-TVB into MOR-expressing cells in the mouse striatum. Scale bar: 200 µm. (D) The number of EnvB-RVΔG-EGFP-infected cells with/without βed-f2-TVB in the mouse striatum. \*\*\**p* < 0.001, vs. βed-f2-TVB at 0 µM (Unpaired *t*-test, n = 3). (E) Targeting efficiency of the bridge protein-mediated EnvB-RVΔG-EGFP infection of MOR-expressing cells in the mouse striatum.

### Bridge protein-mediated viral targeting to MOR-expressing cells in NHPs

In the final set of experiments, we sought to determine whether our βed-f2-TVB bridge protein can effectively target MOR-expressing cells in NHPs. We performed a series of injections into the striatum of a female Japanese macaque under four experimental conditions: EnvB-LV-CAG-EGFP with or without βed-f2-TVB and EnvB-LV-hSyn-tTA-cHS4-tetO-mScarlet3 with or without βed-f2-TVB (Fig. 6A). To assess the specificity of βed-f2-TVB for MOR-positive neurons, we conducted *in situ* hybridization on striatum sections and quantified the colocalization ratio of a fluorescent protein (*EGFP* or *mScarlet3* mRNA) derived from LV vectors with the *MOR* (*Oprm1*) mRNA because the anti-MOR antibody did not perform reliably on the monkey tissue sections. At a concentration of 20 µM βed-f2-TVB, EnvB-LV-CAG-EGFP infected cells around the injection site of the striatum, whereas little *EGFP* expression was detected under bridge protein βed-f2-TVB absence (Fig. 6B, C). This result is consistent with our results in mice (Fig. 4). To determine the specificity of the viral infection for MOR-expressing cells, we performed *in situ* hybridization for *EGFP* and *MOR (Oprm1)*. The percentage of EnvB-LV-CAG-EGFP infected cells co-expressing *EGFP* and *MOR* was 54.1% at 20 µM βed-f2-TVB (Fig. 6D).

**Fig. 6.**
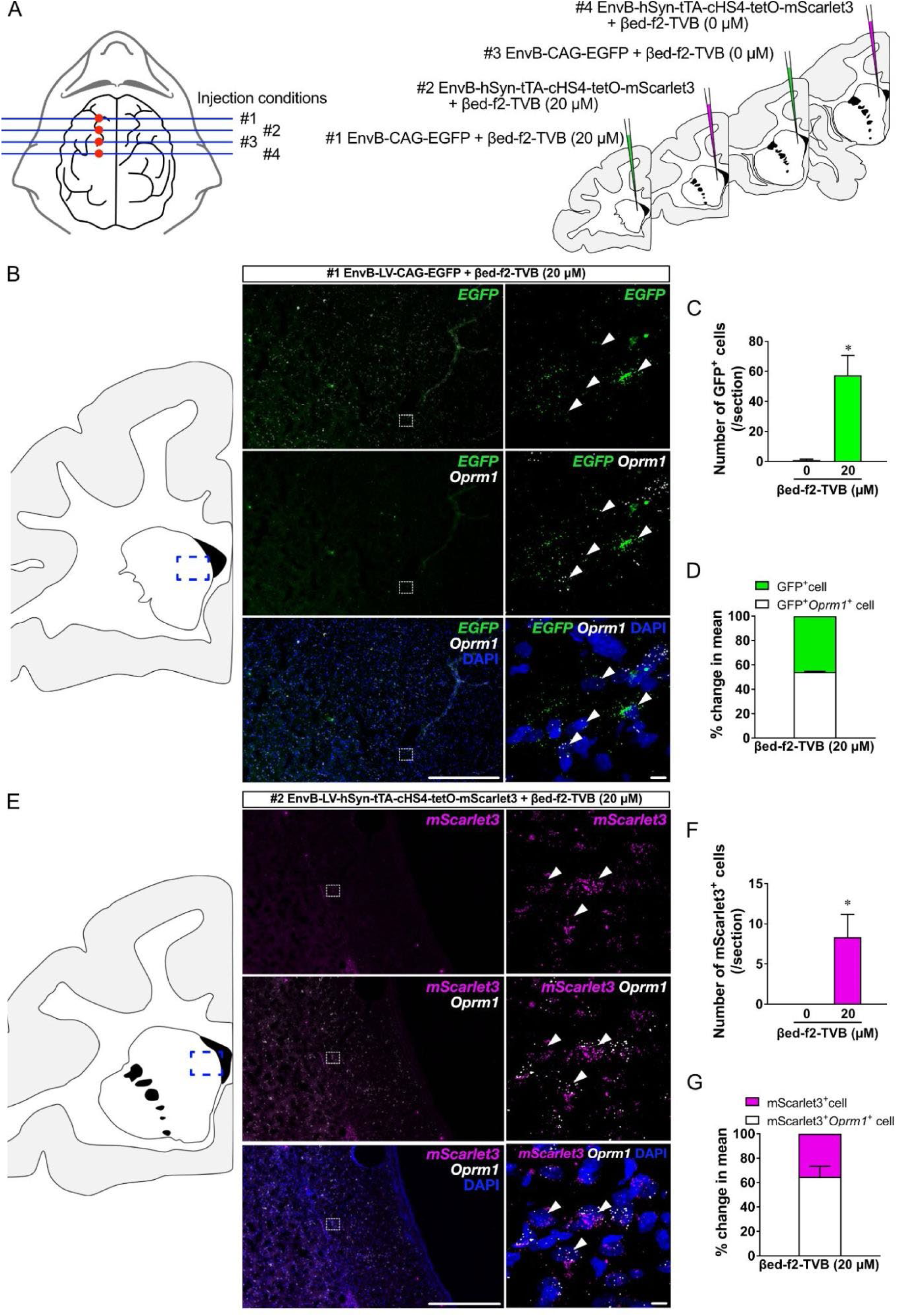
Bridge protein-mediated viral targeting in wild-type monkeys. (A) Experimental conditions of *in vivo* evaluation in monkeys. (Left) Red circles indicate the injection sites along the AP axis of the striatum (#1-4). (Right) #1 and #3 indicate the injection of EnvB-LV-CAG-EGFP with and without βed-f2-TVB, respectively. #2 and #4 indicate the injection of EnvB-LV-hSyn-tTA-cHS4-tetO-mScaret3 with and without βed-f2-TVB, respectively. (B) *In situ* hybridization for *EGFP* and *Oprm* mRNA in brain sections for the injection of EnvB-LV-CAG-EGFP with or without βed-f2-TVB. The left and right panels indicate low- and high-magnification views of the *EGFP*-positive cells in the presence of βed-f2-TVB, respectively. Scale bar: 500 (left) and 20 µm (right panels). (C) The number of EnvB-LV-CAG-EGFP-infected, *EGFP*-positive cells with or without βed-f2-TVB. \**p*<0.05, vs. βed-f2-TVB 0 µM (Unpaired *t*-test, n = 3 sections). (D) *Oprm1*-positive ratio in βed-f2-TVB-mediated EnvB-LV-CAG-EGFP-infected, *EGFP*-positive cells. (E) *In situ* hybridization for *mScarlet3* and *Oprm* mRNA in brain sections for injection of EnvB-LV-hSyn-tTA-cHS4-tetO-mScarlet3 with or without βed-f2-TVB. The left and right panels indicate low- and high-magnification views of the *mScarlet3*-positive cells in the presence of βed-f2-TVB, respectively. Scale bar: 500 (left) and 20 µm (right panels). (F) The number of EnvB-LV-hSyn-tTA-cHS4-tetO-mScarlet3-infected, *mScarlet3*-positive cells with or without βed-f2-TVB. \**p* < 0.05, vs. βed-f2-TVB 0 µM (Unpaired *t*-test, n = 3 sections). (G) *Oprm1*-positive ratio in βed-f2-TVB-mediated EnvB-LV-hSyn-tTA-cHS4-tetO-mScarlet3-infected, *mScarlet3*-positive cells.

To enhance the specificity of viral targeting to MOR-expressing neurons, we employed the intersectional approach using EnvB-LV-hSyn-tTA-cHS4-tetO-mScarlet3 with βed-f2-TVB. EnvB-LV-hSyn-tTA-cHS4-tetO-mScarlet3-infected, *mScarlet3*-positive cells in the striatum at 20 µM βed-f2-TVB, whereas *mScarlet3*-positive cells were not detected under βed-f2-TVB absence (Fig. 6E, F). *In situ* hybridization for *mScarlet3* and *MOR* (*Oprm1*) mRNA revealed that 64.0% of the EnvB-LV-hSyn-tTA-cHS4-tetO-mScarlet3-infected cells co-expressed *mScarlet3* and *MOR* (Fig. 6G). The MOR specificity of the intersectional approach using EnvB-LV-hSyn-tTA-cHS4-tetO-mScarlet3 with βed-f2-TVB exceeded that of the CAG promoter-based system using EnvB-LV-CAG-EGFP (54.1%), consistent with our observations in the mouse striatum. Thus, this intersectional approach combined with neuron-specific promoters and the bridge protein strategy enabled the targeting of viral vectors to MOR-expressing neurons in wild-type NHPs.

## Discussion

The present study demonstrates the development of a viral targeting system using bridge proteins to selectively target endogenous GPCRs, specifically MOR, expressed in neurons of nontransgenic animals. Choi *et al*. utilized a bridge protein containing NRG1, a ligand for the tyrosine kinase-type receptor ErbB4, to direct EnvB-pseudotyped viruses to inhibitory neurons expressing ErbB4 through ligand-mediated ligand-mediated internalization (Choi et al., 2010; Choi and Callaway, 2011). We extended the concept of this bridge protein approach to GPCRs and designed MOR-specific bridge proteins by fusing MOR peptide ligands (e.g., βed, EM1) and viral receptors (e.g., TVB, TVC, TVE) with a two-repeat flexible linker. The MOR ligand-based bridge protein βed-f2-TVB directed LV and RVΔG vectors into MOR-expressing cells in nontransgenic mice and monkeys. The present study established the feasibility of targeting GPCRs by bridge protein-mediated viral infection, broadening the applicability of this system beyond tyrosine kinase receptors. Bridge protein systems will offer a promising strategy for cell-type-specific viral targeting in nontransgenic animals, particularly NHPs, thereby advancing our understanding of cell-type-specific circuit organization and function.

Our findings underscore the critical importance of rational bridge protein design for achieving efficient and specific viral transduction. Despite the feasibility of selective viral infection using bridge proteins (Choi et al., 2010), the detailed configuration of bridge proteins, including the domain arrangement and interdomain linker length, remains underexplored. Machine learning-based models, such as AlphaFold and SignalIP, offer valuable tools for predicting protein structure and signal peptide sequences (Jumper et al., 2021; Teufel et al., 2022). However, predicting the structure, function, and secretion efficiency of artificially designed proteins with non-native secretion signal peptides remains challenging (Brockmeier et al., 2006; Freudl, 2018). Therefore, experimentally testing various signal peptides, linkers, and domain arrangements is essential to identify effective engineered bridge proteins for viral infection. To the best of our knowledge, the present study is the first to comprehensively demonstrate the effects of signal peptide sequence, ligand orientation, interdomain linker length, and reaction conditions on bridge protein-mediated viral infection. Among these parameters, selecting an appropriate secretion signal peptide is essential for maximizing the stability and bioactivity of recombinant proteins (Freudl, 2018). TVB-based bridge proteins lacking a signal peptide and those incorporating a non-native signal peptide derived from IL-6 or IFNγ failed to induce efficient bridge protein production or viral infection (Fig. 1), and TVC- and TVE-based bridge proteins using a non-native signal peptide derived from TVB were ineffective in viral infection (Fig. 3). Collectively, these results suggest that native TVX-derived signal peptides are suited for use in their corresponding TVX-based bridge proteins, likely due to their favorable effects on protein folding and stability (Freudl, 2018). Additionally, we found that the placement of the MOR ligand at the N terminus of the bridge protein was crucial for effective viral infection (Fig. 2, 3). This result is consistent with prior studies showing that the binding site of the MOR ligand is located at its N terminus (Amiche et al., 1990). Notably, the TVB at the C terminus did not interfere with the binding of the MOR to the bridge protein (Gaudriault et al., 1997), highlighting the importance of ligand orientation in bridge protein design for efficient viral targeting. Furthermore, the interdomain linker length also emerged as a key design parameter (Fig. 2). GGGGS linkers shorter than two repeats can efficiently produce proteins but may cause interference between the MOR ligand and viral receptor domains, whereas excessively long linkers are likely to decrease bridge protein production and stability. These results are consistent with previous studies showing that an appropriate linker length improved the activity of fusion proteins compared to those without linkers or with longer linkers (Bai and Shen 2006). Thus, selecting an appropriate linker length minimized the interference between the MOR ligand and the viral receptor domains while maintaining their activity. Finally, optimizing the bridge protein−virus interaction conditions, such as the bridge protein/virus ratio and incubation time, further enhanced the viral infection efficiency, underscoring the importance of fine-tuning the experimental parameters for bridge protein-mediated viral targeting. Overall, our study identifies critical parameters for the structural design of bridge proteins, including the selection of secretion signal peptide, ligand orientation, and types and lengths of interdomain linkers, and the optimization of reaction conditions. The present study lays the groundwork for the rational design of bridge proteins, which will facilitate the development of bridge proteins for various receptor-ligand pairs.

We estimated the ligand activity of the bridge proteins. Viral infection mediated by βed-f2-TVB and EM1-f2-TVB in CHO-mMOR cells was competitively blocked by the MOR antagonist NOX (Fig. 2K, 3F). These inhibition data were analyzed using the Cheng-Prusoff equation [Ki = IC_50_ / (1 + [L]/Kd)], where Ki represents the inhibitory constant of the antagonist; IC_50_ is the concentration at which the inhibitory ligand displaces 50% of the ligand; [L] is the concentration of the ligand in the assay; and Kd is the affinity constant of the ligand (Cheng and Prusoff, 1973). Here, we defined the ligand activity of bridge proteins as the viral infection activity inducing GFP-positive cells via MORs in CHO-mMOR cells. The MOR antagonist NOX inhibited the ligand activity of βed-f2-TVB and EM1-f2-TVB in a concentration-dependent manner (Fig. 2K, 3F), yielding IC_50_ values. For βed-f2-TVB (Fig. 2K), the IC_50_ of NOX was determined to be 2.3 mM. Given the reported Kd for βed of 1.8 nM (Matsui et al., 1985) and the Ki for NOX of 1.4 nM (Romero et al., 1999), the concentration of the ligand activity of the virus-bound βed-f2-TVB was calculated to be approximately 3.0 mM using the Cheng-Prusoff equation. Similarly, for EM1-f2-TVB (3F), the IC_50_ of NOX was determined to be 1.5 mM. Using the Kd for EM1 of 1.1 nM (Hosohata et al., 1998) and the Ki for NOX of 1.4 nM (Romero et al., 1999), the ligand activity concentration of virus-bound EM1-f2-TVB was calculated to be 1.2 mM using the Cheng-Prusoff equation. The difference in ligand activity concentrations between virus-bound βed-f2-TVB (2.0 mM) and virus-bound EM1-f2-TVB (1.2 mM) with different Kd values suggests that higher concentrations of EM1-f2-TVB are required to achieve the same level of viral infection (∼90 GFP-expressing cells/well) as βed-f2-TVB in CHO-mMOR cells. Conceivably, bridge proteins using ligands with lower Kd values, indicative of higher affinity with target receptors, provide higher viral infection efficiency when using the equivalent amounts of viral vectors and bridge proteins. Thus, the Kd value of ligands is considered a critical factor in the rational design of bridge proteins, with lower Kd values increasing infection efficiency.

The striatum serves as a gateway to the basal ganglia and plays a pivotal role in motor planning and action selection (Graybiel and Matsushima, 2023; Klaus et al., 2019). Striatum dysfunction has been implicated in a wide range of neurological and psychiatric disorders, such as Parkinson’s disease, Huntington’s disease, Tourette’s syndrome, schizophrenia, and addiction (Kreitzer and Malenka, 2008; Nambu et al., 2023). The striatum comprises heterogeneous cell types in distinct compartments, and information processing in the striatum depends on cell-type-specific circuits. The distinct roles of dopamine D1 receptor (D1R)^+^ cells and dopamine D2 receptor (D2R)^+^ cells, and the striosome and matrix compartments have been studied (Matsushima et al., 2023). Most studies have mainly focused on the D1R^+^ direct vs. D2R^+^ indirect pathways by leveraging the availability of transgenic animals that label D1R- or D2R-expressing cells. However, recent single-cell transcriptome analyses revealed heterogeneity within the D1R- and D2R-positive medium spiny neurons (Gokce et al., 2016; Stanley et al., 2020), suggesting that the traditional model that involves competition between signals in the D1R^+^ direct and D2R^+^ indirect pathways does not fully account for the broad functional roles of the striatum. These studies highlight the need for higher-resolution tools to dissect cell types and their functional roles in the striatum. In contrast, the function and input–output relationship of the compartmental structure of the striatum remain poorly understood. Studies of the striosome (also known as the patch) and matrix compartments of the striatum have been limited, primarily due to the methodological challenges in selectively targeting cells in these compartments (Graybiel and Matsushima, 2023). In rodents, MOR-expressing neurons in the striatum largely correspond to the striosome. However, the degree of overlap between MOR expression and the striosome is relatively lower in NHPs (He et al., 2021). These findings suggest that MOR-expressing neurons in the primate striatum have a unique spatial and compartmental organization, highlighting the need to investigate this structure specifically in NHPs.

Targeting specific neuronal cell types in the primate striatum using viral vectors remains a notable challenge (Salimando et al., 2023). In this study, we demonstrated that viral targeting via bridge proteins can be applied to MOR-expressing neurons in the striatum of nontransgenic mice and monkeys, with a moderate specificity of approximately 54%–65%. The βed-f2-TVB-mediated viral infection mediates the specific interaction between the bridge protein and MORs on the target cells, supported by three findings: (i) EnvB-pseudotyped virus vectors under bridge protein absence did not express reporter genes; (ii) βed-f2-TVB-mediated viral infection was observed in CHO-mMOR cells, but not in CHO cells; and (iii) the MOR antagonist NOX competitively inhibited βed-f2-TVB-mediated viral infection. However, some off-target viral infections were observed. This could potentially be attributed to the affinity of the MOR ligands βed and EM1 for other receptors, such as κ-opioid receptors (KORs) and δ-opioid receptors (DORs). This non-specific binding could lead to viral infections in MOR-negative but KOR/DOR-positive cells. Addressing this cross-reactivity is crucial for enhancing targeting specificity. Nontransgenic approaches for targeting specific cell types have garnered considerable interest. For example, cell type-specific *cis*-regulatory elements, such as promoters and enhancers, have been identified and incorporated into AAV vectors (Graybuck et al., 2021; Jüttner et al., 2019; Mich et al., 2021; Salimando et al., 2023). AAV-based systems called CellREADR and READR utilized endogenous mRNA-dependent translation of exogenous mRNA (Kaseniit et al., 2023; Qian et al., 2022). Lentivirus vectors displaying engineered viral envelopes with particular ligands infected cells expressing receptors for displayed ligands (Hughes et al., 2024; Neda et al., 1991; Strebinger et al., 2023; Yu et al., 2022). Despite these advances, no single viral approach has yet achieved ∼100% cell-type specificity. This limitation highlights the need for intersectional strategies that combine multiple molecular approaches to further enhance targeting specificity. Using combinations of recombinase, such as Cre, Flp, and Dre, and tTA, will also facilitate intersectional strategies (Fenno et al., 2020, 2014; Hughes et al., 2024). Overall, integrating various molecular approaches with the bridge protein system could enhance the precision of targeting cell populations.

The present study extends the application of bridge proteins to other ASLV envelope subtypes/receptor systems, such as the EnvC/TVC and EnvE/TVE systems, analogous to the EnvB/TVB system. Multiplex viral targeting of three independent cell populations can be achieved with bridge proteins using these three distinct EnvX/TVX systems within the same nontransgenic animals, including NHPs (Osakada and Callaway, 2013; Suzuki et al., 2020). In addition, we have shown that bridge protein systems can target not only tyrosine kinase receptors but also GPCRs on specific cell types using enveloped viruses. These findings underscore the versatility of bridge proteins and their potential for broader applications in revealing cell-type-specific functions in complex biological systems. Future research should focus on extending the target range of bridge proteins to other classes of surface proteins, including ion channels and transporters. A key next step is to investigate the use of small molecule ligands for the bridge protein-mediated targeting of viral infection, as many specific pharmacological chemical ligands are already available. Developing a bridge protein library containing ligands for various surface receptors could greatly expand the range of targetable cell types, not only within the brain but also across the whole body (Bian et al., 2005).

## Author contributions

R.K. performed the molecular cloning, bridge protein production, virus production, infection assay, virus injection, and histological analysis, collected and analyzed all data, and wrote the manuscript. S.A. and K.A. performed the monkey surgery, virus injection, and histological analysis. F.O. wrote the manuscript and supervised the project.

## Declaration of Competing Interest

The authors have no competing interests to declare.

## Acknowledgements

We thank all members of the Osakada laboratory for their discussions and Jung-min OH (Kyoto University) for the monkey surgery.

## Fundings

This work was supported by the Nagoya University Interdisciplinary Frontier Fellowship (R.K.), JSPS (S.A., 21J40030 and 21K07259; K.A., 20H03555; and F.O., 18H02706, 20K21476, and 21H05168), AMED (F.O., JP20dm0207058h, JP23gm1510011h, and JP24wm0625110h), JST (F.O., JPMJPR14F6 and JPMJCR1851), ASHBi flagship project (K.A.), Takeda Science Foundation (S.A. and F.O.), Naito Foundation (S.A.), SRF Foundation (F.O.), and Suzuken Memorial Foundation (F.O.).

## Notes

### Competing Interest Statement

The authors have declared no competing interest.

## References

Adamczyk, A., 2023. Glial-neuronal interactions in neurological disorders: Molecular mechanisms and potential points for intervention. Int. J. Mol. Sci. 24, 6274.

Alfthan, K., Takkinen, K., Sizmann, D., Soderlund, H., Teeri, T.T., 1995. Properties of a single-chain antibody containing different linker peptides. Protein Eng. Des. Sel. 8, 725–731.

Amiche, M., Sagan, S., Mor, A., Pelaprat, D., Rostene, W., Delfour, A., Nicolas, P., 1990. Characterisation and visualisation of [3H]dermorphin binding to mu opioid receptors in the rat brain. Combined high selectivity and affinity in a natural peptide agonist for the morphine (mu) receptor. Eur. J. Biochem. 189, 625–635.

Arai, R., Ueda, H., Kitayama, A., Kamiya, N., Nagamune, T., 2001. Design of the linkers which effectively separate domains of a bifunctional fusion protein. Protein Eng. 14, 529–532.

Argos, P., 1990. An investigation of oligopeptides linking domains in protein tertiary structures and possible candidates for general gene fusion. J. Mol. Biol. 211, 943–958.

Arttamangkul, S., Plazek, A., Platt, E.J., Jin, H., Murray, T.F., Birdsong, W.T., Rice, K.C., Farrens, D.L., Williams, J.T., 2019. Visualizing endogenous opioid receptors in living neurons using ligand-directed chemistry. Elife 8. 10.7554/eLife.49319

Atasoy, D., Aponte, Y., Su, H.H., Sternson, S.M., 2008. A FLEX switch targets Channelrhodopsin-2 to multiple cell types for imaging and long-range circuit mapping. J. Neurosci. 28, 7025–7030.

Bai, Y., Shen, W.-C., 2006. Improving the oral efficacy of recombinant granulocyte colony-stimulating factor and transferrin fusion protein by spacer optimization. Pharm. Res. 23, 2116–2121.

Balliet, J.W., Berson, J., D’Cruz, C.M., Huang, J., Crane, J., Gilbert, J.M., Bates, P., 1999. Production and characterization of a soluble, active form of Tva, the subgroup A avian sarcoma and leukosis virus receptor. J. Virol. 73, 3054–3061.

Barnard, R.J.O., Elleder, D., Young, J.A.T., 2006. Avian sarcoma and leukosis virus-receptor interactions: from classical genetics to novel insights into virus-cell membrane fusion. Virology 344, 25–29.

Barnard, R.J.O., Young, J.A.T., 2003. Alpharetrovirus envelope-receptor interactions. Curr. Top. Microbiol. Immunol. 281, 107–136.

Bian, H., Fournier, P., Moormann, R., Peeters, B., Schirrmacher, V., 2005. Selective gene transfer to tumor cells by recombinant Newcastle Disease Virus via a bispecific fusion protein. Int. J. Oncol. 26, 431–439.

Boerger, A.L., Snitkovsky, S., Young, J.A.T., 1999. Retroviral vectors preloaded with a viral receptor-ligand bridge protein are targeted to specific cell types. Proc. Natl. Acad. Sci. U. S. A. 96, 9867–9872.

Branda, C.S., Dymecki, S.M., 2004. Talking about a revolution: The impact of site-specific recombinases on genetic analyses in mice. Dev. Cell 6, 7–28.

Brimblecombe, K.R., Cragg, S.J., 2017. The Striosome and Matrix Compartments of the Striatum: A Path through the Labyrinth from Neurochemistry toward Function. ACS Chem. Neurosci. 8, 235–242.

Brockmeier, U., Caspers, M., Freudl, R., Jockwer, A., Noll, T., Eggert, T., 2006. Systematic screening of all signal peptides from Bacillus subtilis: a powerful strategy in optimizing heterologous protein secretion in Gram-positive bacteria. J. Mol. Biol. 362, 393–402.

Cheng, Y., Prusoff, W.H., 1973. Relationship between the inhibition constant (K1) and the concentration of inhibitor which causes 50 per cent inhibition (I50) of an enzymatic reaction. Biochem. Pharmacol. 22, 3099–3108.

Chen, X., Ravindra Kumar, S., Adams, C.D., Yang, D., Wang, T., Wolfe, D.A., Arokiaraj, C.M., Ngo, V., Campos, L.J., Griffiths, J.A., Ichiki, T., Mazmanian, S.K., Osborne, P.B., Keast, J.R., Miller, C.T., Fox, A.S., Chiu, I.M., Gradinaru, V., 2022. Engineered AAVs for non-invasive gene delivery to rodent and non-human primate nervous systems. Neuron 110, 2242–2257.e6.

Chen, X., Zaro, J.L., Shen, W.-C., 2013. Fusion protein linkers: property, design and functionality. Adv. Drug Deliv. Rev. 65, 1357–1369.

Choi, J., Callaway, E.M., 2011. Monosynaptic inputs to ErbB4-expressing inhibitory neurons in mouse primary somatosensory cortex. J. Comp. Neurol. 519, 3402–3414.

Choi, J., Young, J.A.T., Callaway, E.M., 2010. Selective viral vector transduction of ErbB4 expressing cortical interneurons in vivo with a viral receptor-ligand bridge protein. Proc. Natl. Acad. Sci. U. S. A. 107, 16703–16708.

Chuapoco, M.R., Flytzanis, N.C., Goeden, N., Christopher Octeau, J., Roxas, K.M., Chan, K.Y., Scherrer, J., Winchester, J., Blackburn, R.J., Campos, L.J., Man, K.N.M., Sun, J., Chen, X., Lefevre, A., Singh, V.P., Arokiaraj, C.M., Shay, T.F., Vendemiatti, J., Jang, M.J., Mich, J.K., Bishaw, Y., Gore, B.B., Omstead, V., Taskin, N., Weed, N., Levi, B.P., Ting, J.T., Miller, C.T., Deverman, B.E., Pickel, J., Tian, L., Fox, A.S., Gradinaru, V., 2023. Adeno-associated viral vectors for functional intravenous gene transfer throughout the non-human primate brain. Nat. Nanotechnol. 18, 1241–1251.

Delfs, J.M., Kong, H., Mestek, A., Chen, Y., Yu, L., Reisine, T., Chesselet, M.F., 1994. Expression of mu opioid receptor mRNA in rat brain: an in situ hybridization study at the single cell level. J. Comp. Neurol. 345, 46–68.

Ding, S., Wu, X., Li, G., Han, M., Zhuang, Y., Xu, T., 2005. Efficient transposition of the piggyBac (PB) transposon in mammalian cells and mice. Cell 122, 473–483.

Dobson, C.S., Reich, A.N., Gaglione, S., Smith, B.E., Kim, E.J., Dong, J., Ronsard, L., Okonkwo, V., Lingwood, D., Dougan, M., Dougan, S.K., Birnbaum, M.E., 2022. Antigen identification and high-throughput interaction mapping by reprogramming viral entry. Nat. Methods 19, 449–460.

Erbs, E., Faget, L., Scherrer, G., Matifas, A., Filliol, D., Vonesch, J.-L., Koch, M., Kessler, P., Hentsch, D., Birling, M.-C., Koutsourakis, M., Vasseur, L., Veinante, P., Kieffer, B.L., Massotte, D., 2015. A mu–delta opioid receptor brain atlas reveals neuronal co-occurrence in subcortical networks. Brain Struct. Funct. 220, 677–702.

Fenno, L.E., Mattis, J., Ramakrishnan, C., Hyun, M., Lee, S.Y., He, M., Tucciarone, J., Selimbeyoglu, A., Berndt, A., Grosenick, L., Zalocusky, K.A., Bernstein, H., Swanson, H., Perry, C., Diester, I., Boyce, F.M., Bass, C.E., Neve, R., Huang, Z.J., Deisseroth, K., 2014. Targeting cells with single vectors using multiple-feature Boolean logic. Nat. Methods 11, 763–772.

Fenno, L.E., Ramakrishnan, C., Kim, Y.S., Evans, K.E., Lo, M., Vesuna, S., Inoue, M., Cheung, K.Y.M., Yuen, E., Pichamoorthy, N., Hong, A.S.O., Deisseroth, K., 2020. Comprehensive dual- and triple-feature intersectional single-vector delivery of diverse functional payloads to cells of behaving mammals. Neuron 107, 836–853.e11.

Freudl, R., 2018. Signal peptides for recombinant protein secretion in bacterial expression systems. Microb. Cell Fact. 17, 52.

Gaudriault, G., Nouel, D., Farra, C.D., Beaudet, A., Vincent, J.-P., 1997. Receptor-induced Internalization of Selective Peptidic μ and Δ Opioid Ligands*. J. Biol. Chem. 272, 2880–2888.

Gokce, O., Stanley, G.M., Treutlein, B., Neff, N.F., Camp, J.G., Malenka, R.C., Rothwell, P.E., Fuccillo, M.V., Südhof, T.C., Quake, S.R., 2016. Cellular taxonomy of the mouse striatum as revealed by single-cell RNA-seq. Cell Rep. 16, 1126–1137.

Graybiel, A.M., Matsushima, A., 2023. Striosomes and Matrisomes: Scaffolds for Dynamic Coupling of Volition and Action. Annu. Rev. Neurosci. 10.1146/annurev-neuro-121522-025740

Graybuck, L.T., Daigle, T.L., Sedeño-Cortés, A.E., Walker, M., Kalmbach, B., Lenz, G.H., Morin, E., Nguyen, T.N., Garren, E., Bendrick, J.L., Kim, T.K., Zhou, T., Mortrud, M., Yao, S., Siverts, L.A., Larsen, R., Gore, B.B., Szelenyi, E.R., Trader, C., Balaram, P., van Velthoven, C.T.J., Chiang, M., Mich, J.K., Dee, N., Goldy, J., Cetin, A.H., Smith, K., Way, S.W., Esposito, L., Yao, Z., Gradinaru, V., Sunkin, S.M., Lein, E., Levi, B.P., Ting, J.T., Zeng, H., Tasic, B., 2021. Enhancer viruses for combinatorial cell-subclass-specific labeling. Neuron 109, 1449–1464.e13.

Han, Y., Sun, X., Kuang, D., Tong, P., Lu, C., Wang, W., Li, N., Chen, Y., Wang, X., Dai, J., Zhang, H., 2017. Characterization of tree shrew (Tupaia belangeri) interleukin-6 and its expression pattern in response to exogenous challenge. Int. J. Mol. Med. 40, 1679–1690.

He, J., Kleyman, M., Chen, J., Alikaya, A., Rothenhoefer, K.M., Ozturk, B.E., Wirthlin, M., Bostan, A.C., Fish, K., Byrne, L.C., Pfenning, A.R., Stauffer, W.R., 2021. Transcriptional and anatomical diversity of medium spiny neurons in the primate striatum. Curr. Biol. 31, 5473–5486.e6.

Hioki, H., Kuramoto, E., Konno, M., Kameda, H., Takahashi, Y., Nakano, T., Nakamura, K.C., Kaneko, T., 2009. High-level transgene expression in neurons by lentivirus with Tet-Off system. Neurosci. Res. 63, 149–154.

Horii, T., Morita, S., Kimura, M., Terawaki, N., Shibutani, M., Hatada, I., 2017. Efficient generation of conditional knockout mice via sequential introduction of lox sites. Sci. Rep. 7, 7891.

Hosohata, K., Burkey, T.H., Alfaro-Lopez, J., Varga, E., Hruby, V.J., Roeske, W.R., Yamamura, H.I., 1998. Endomorphin-1 and endomorphin-2 are partial agonists at the human mu-opioid receptor. Eur. J. Pharmacol. 346, 111–114.

Huang, Z.J., Zeng, H., 2013. Genetic approaches to neural circuits in the mouse. Annu. Rev. Neurosci. 36, 183–215.

Hughes, A.C., Pittman, B.G., Xu, B., Gammons, J.W., Webb, C.M., Nolen, H.G., Chapman, P., Bikoff, J.B., Schwarz, L.A., 2024. A single-vector intersectional AAV strategy for interrogating cellular diversity and brain function. Nat. Neurosci. 27, 1400–1410.

Johnston, J.G., Gerfen, C.R., Haber, S.N., van der Kooy, D., 1990. Mechanisms of striatal pattern formation: conservation of mammalian compartmentalization. Brain Res. Dev. Brain Res. 57, 93–102.

Jumper, J., Evans, R., Pritzel, A., Green, T., Figurnov, M., Ronneberger, O., Tunyasuvunakool, K., Bates, R., Žídek, A., Potapenko, A., Bridgland, A., Meyer, C., Kohl, S.A.A., Ballard, A.J., Cowie, A., Romera-Paredes, B., Nikolov, S., Jain, R., Adler, J., Back, T., Petersen, S., Reiman, D., Clancy, E., Zielinski, M., Steinegger, M., Pacholska, M., Berghammer, T., Bodenstein, S., Silver, D., Vinyals, O., Senior, A.W., Kavukcuoglu, K., Kohli, P., Hassabis, D., 2021. Highly accurate protein structure prediction with AlphaFold. Nature 596, 583–589.

Jüttner, J., Szabo, A., Gross-Scherf, B., Morikawa, R.K., Rompani, S.B., Hantz, P., Szikra, T., Esposti, F., Cowan, C.S., Bharioke, A., Patino-Alvarez, C.P., Keles, Ö., Kusnyerik, A., Azoulay, T., Hartl, D., Krebs, A.R., Schübeler, D., Hajdu, R.I., Lukats, A., Nemeth, J., Nagy, Z.Z., Wu, K.-C., Wu, R.-H., Xiang, L., Fang, X.-L., Jin, Z.-B., Goldblum, D., Hasler, P.W., Scholl, H.P.N., Krol, J., Roska, B., 2019. Targeting neuronal and glial cell types with synthetic promoter AAVs in mice, non-human primates and humans. Nat. Neurosci. 22, 1345–1356.

Kaseniit, K.E., Katz, N., Kolber, N.S., Call, C.C., Wengier, D.L., Cody, W.B., Sattely, E.S., Gao, X.J., 2023. Modular, programmable RNA sensing using ADAR editing in living cells. Nat. Biotechnol. 41, 482–487.

Kishi, N., Sato, K., Sasaki, E., Okano, H., 2014. Common marmoset as a new model animal for neuroscience research and genome editing technology. Dev. Growth Differ. 56, 53–62.

Klaus, A., Alves da Silva, J., Costa, R.M., 2019. What, if, and when to move: Basal ganglia circuits and self-paced action initiation. Annu. Rev. Neurosci. 42, 459–483.

Kodera, T., Takeuchi, R.F., Takahashi, S., Suzuki, K., Kassai, H., Aiba, A., Shiozawa, S., Okano, H., Osakada, F., 2023. Modeling the marmoset brain using embryonic stem cell-derived cerebral assembloids. Biochem. Biophys. Res. Commun. 657, 119–127.

Kreitzer, A.C., Malenka, R.C., 2008. Striatal plasticity and basal ganglia circuit function. Neuron 60, 543–554.

Luo, L., 2021. Architectures of neuronal circuits. Science 373, eabg7285.

Luo, L., Callaway, E.M., Svoboda, K., 2018. Genetic Dissection of Neural Circuits: A Decade of Progress. Neuron 98, 865.

Maduna, T., Audouard, E., Dembélé, D., Mouzaoui, N., Reiss, D., Massotte, D., Gaveriaux-Ruff, C., 2018. Microglia Express Mu Opioid Receptor: Insights From Transcriptomics and Fluorescent Reporter Mice. Front. Psychiatry 9, 726.

Mansour, A., Fox, C.A., Burke, S., Meng, F., Thompson, R.C., Akil, H., Watson, S.J., 1994. Mu, delta, and kappa opioid receptor mRNA expression in the rat CNS: an in situ hybridization study. J. Comp. Neurol. 350, 412–438.

Masaki, Y., Yamaguchi, M., Takeuchi, R.F., Osakada, F., 2022. Monosynaptic rabies virus tracing from projection-targeted single neurons. Neurosci. Res. 178, 20–32.

Matsui, H., Asakura, M., Tsukamoto, T., Imafuku, J., Ino, M., Saitoh, N., Miyamura, S., Hasegawa, K., 1985. Solubilization and characterization of rat brain alpha 2-adrenergic receptor. J. Neurochem. 44, 1625–1632.

Matsushima, A., Pineda, S.S., Crittenden, J.R., Lee, H., Galani, K., Mantero, J., Tombaugh, G., Kellis, M., Heiman, M., Graybiel, A.M., 2023. Transcriptional vulnerabilities of striatal neurons in human and rodent models of Huntington’s disease. Nat. Commun. 14, 1–17.

Matsuyama, M., Ohashi, Y., Tsubota, T., Yaguchi, M., Kato, S., Kobayashi, K., Miyashita, Y., 2015. Avian sarcoma leukosis virus receptor-envelope system for simultaneous dissection of multiple neural circuits in mammalian brain. Proc. Natl. Acad. Sci. U. S. A. 112, E2947–E2956.

McConalogue, K., Grady, E.F., Minnis, J., Balestra, B., Tonini, M., Brecha, N.C., Bunnett, N.W., Sternini, C., 1999. Activation and internalization of the mu-opioid receptor by the newly discovered endogenous agonists, endomorphin-1 and endomorphin-2. Neuroscience 90, 1051–1059.

Mich, J.K., Graybuck, L.T., Hess, E.E., Mahoney, J.T., Kojima, Y., Ding, Y., Somasundaram, S., Miller, J.A., Kalmbach, B.E., Radaelli, C., Gore, B.B., Weed, N., Omstead, V., Bishaw, Y., Shapovalova, N.V., Martinez, R.A., Fong, O., Yao, S., Mortrud, M., Chong, P., Loftus, L., Bertagnolli, D., Goldy, J., Casper, T., Dee, N., Opitz-Araya, X., Cetin, A., Smith, K.A., Gwinn, R.P., Cobbs, C., Ko, A.L., Ojemann, J.G., Keene, C.D., Silbergeld, D.L., Sunkin, S.M., Gradinaru, V., Horwitz, G.D., Zeng, H., Tasic, B., Lein, E.S., Ting, J.T., Levi, B.P., 2021. Functional enhancer elements drive subclass-selective expression from mouse to primate neocortex. Cell Rep. 34, 108754.

Minami, M., Satoh, M., 1995. Molecular biology of the opioid receptors: structures, functions and distributions 23, 121–145.

Mores, K.L., Cassell, R.J., van Rijn, R.M., 2019. Arrestin recruitment and signaling by G protein-coupled receptor heteromers. Neuropharmacology 152, 15–21.

Mothes, W., Boerger, A.L., Narayan, S., Cunningham, J.M., Young, J.A., 2000. Retroviral entry mediated by receptor priming and low pH triggering of an envelope glycoprotein. Cell 103, 679–689.

Nagai, J., Yu, X., Papouin, T., Cheong, E., Freeman, M.R., Monk, K.R., Hastings, M.H., Haydon, P.G., Rowitch, D., Shaham, S., Khakh, B.S., 2021. Behaviorally consequential astrocytic regulation of neural circuits. Neuron 109, 576–596.

Nambu, A., Chiken, S., Sano, H., Hatanaka, N., Obeso, J.A., 2023. Dynamic activity model of movement disorders: The fundamental role of the hyperdirect pathway. Mov. Disord. 38, 2145–2150.

Nectow, A.R., Nestler, E.J., 2020. Viral tools for neuroscience. Nat. Rev. Neurosci. 21, 669–681.

Neda, H., Wu, C.H., Wu, G.Y., 1991. Chemical modification of an ecotropic murine leukemia virus results in redirection of its target cell specificity. J. Biol. Chem. 266, 14143–14146.

Ohtsuka, M., Sato, M., Miura, H., Takabayashi, S., Matsuyama, M., Koyano, T., Arifin, N., Nakamura, S., Wada, K., Gurumurthy, C.B., 2018. i-GONAD: a robust method for in situ germline genome engineering using CRISPR nucleases. Genome Biol. 19, 25.

Okigawa, S., Yamaguchi, M., Ito, K.N., Takeuchi, R.F., Morimoto, N., Osakada, F., 2021. Cell type- and layer-specific convergence in core and shell neurons of the dorsal lateral geniculate nucleus. J. Comp. Neurol. 529, 2099–2124.

Osakada, F., Callaway, E.M., 2013. Design and generation of recombinant rabies virus vectors. Nat. Protoc. 8, 1583–1601.

Osakada, F., Mori, T., Cetin, A.H., Marshel, J.H., Virgen, B., Callaway, E.M., 2011. New rabies virus variants for monitoring and manipulating activity and gene expression in defined neural circuits. Neuron 71, 617–631.

Pan, Y.C., Stern, A.S., Familletti, P.C., Khan, F.R., Chizzonite, R., 1987. Structural characterization of human interferon gamma. Heterogeneity of the carboxyl terminus. Eur. J. Biochem. 166, 145–149.

Paolicelli, R.C., Sierra, A., Stevens, B., Tremblay, M.-E., Aguzzi, A., Ajami, B., Amit, I., Audinat, E., Bechmann, I., Bennett, M., Bennett, F., Bessis, A., Biber, K., Bilbo, S., Blurton-Jones, M., Boddeke, E., Brites, D., Brône, B., Brown, G.C., Butovsky, O., Carson, M.J., Castellano, B., Colonna, M., Cowley, S.A., Cunningham, C., Davalos, D., De Jager, P.L., de Strooper, B., Denes, A., Eggen, B.J.L., Eyo, U., Galea, E., Garel, S., Ginhoux, F., Glass, C.K., Gokce, O., Gomez-Nicola, D., González, B., Gordon, S., Graeber, M.B., Greenhalgh, A.D., Gressens, P., Greter, M., Gutmann, D.H., Haass, C., Heneka, M.T., Heppner, F.L., Hong, S., Hume, D.A., Jung, S., Kettenmann, H., Kipnis, J., Koyama, R., Lemke, G., Lynch, M., Majewska, A., Malcangio, M., Malm, T., Mancuso, R., Masuda, T., Matteoli, M., McColl, B.W., Miron, V.E., Molofsky, A.V., Monje, M., Mracsko, E., Nadjar, A., Neher, J.J., Neniskyte, U., Neumann, H., Noda, M., Peng, B., Peri, F., Perry, V.H., Popovich, P.G., Pridans, C., Priller, J., Prinz, M., Ragozzino, D., Ransohoff, R.M., Salter, M.W., Schaefer, A., Schafer, D.P., Schwartz, M., Simons, M., Smith, C.J., Streit, W.J., Tay, T.L., Tsai, L.-H., Verkhratsky, A., von Bernhardi, R., Wake, H., Wittamer, V., Wolf, S.A., Wu, L.-J., Wyss-Coray, T., 2022. Microglia states and nomenclature: A field at its crossroads. Neuron 110, 3458–3483.

Park, J.E., Silva, A.C., 2019. Generation of genetically engineered non-human primate models of brain function and neurological disorders. Am. J. Primatol. 81, e22931.

Qian, Y., Li, J., Zhao, S., Matthews, E.A., Adoff, M., Zhong, W., An, X., Yeo, M., Park, C., Yang, X., Wang, B.-S., Southwell, D.G., Huang, Z.J., 2022. Programmable RNA sensing for cell monitoring and manipulation. Nature 610, 713–721.

Reinisová, M., Senigl, F., Yin, X., Plachy, J., Geryk, J., Elleder, D., Svoboda, J., Federspiel, M.J., Hejnar, J., 2008. A single-amino-acid substitution in the TvbS1 receptor results in decreased susceptibility to infection by avian sarcoma and leukosis virus subgroups B and D and resistance to infection by subgroup E in vitro and in vivo. J. Virol. 82, 2097–2105.

Rinderknecht, E., O’Connor, B.H., Rodriguez, H., 1984. Natural human interferon-gamma. Complete amino acid sequence and determination of sites of glycosylation. J. Biol. Chem. 259, 6790–6797.

Romero, D.V., Partilla, J.S., Zheng, Q.X., Heyliger, S.O., Ni, Q., Rice, K.C., Lai, J., Rothman, R.B., 1999. Opioid peptide receptor studies. 12. Buprenorphine is a potent and selective mu/kappa antagonist in the [35S]-GTP-gamma-S functional binding assay. Synapse 34, 83–94.

Salimando, G.J., Tremblay, S., Kimmey, B.A., Li, J., Rogers, S.A., Wojick, J.A., McCall, N.M., Wooldridge, L.M., Rodrigues, A., Borner, T., Gardiner, K.L., Jayakar, S.S., Singeç, I., Woolf, C.J., Hayes, M.R., De Jonghe, B.C., Bennett, F.C., Bennett, M.L., Blendy, J.A., Platt, M.L., Creasy, K.T., Renthal, W.R., Ramakrishnan, C., Deisseroth, K., Corder, G., 2023. Human OPRM1 and murine Oprm1 promoter driven viral constructs for genetic access to μ-opioidergic cell types. Nat. Commun. 14, 5632.

Shinmyo, Y., Terashita, Y., Dinh Duong, T.A., Horiike, T., Kawasumi, M., Hosomichi, K., Tajima, A., Kawasaki, H., 2017. Folding of the cerebral cortex requires Cdk5 in upper-layer neurons in gyrencephalic mammals. Cell Rep. 20, 2131–2143.

Smith, J.G., Mothes, W., Blacklow, S.C., Cunningham, J.M., 2004. The mature avian leukosis virus subgroup A envelope glycoprotein is metastable, and refolding induced by the synergistic effects of receptor binding and low pH is coupled to infection. J. Virol. 78, 1403–1410.

Snitkovsky, S., Niederman, T.M.J., Mulligan, R.C., Young, J.A.T., 2001. Targeting Avian Leukosis Virus Subgroup A Vectors by Using a TVA-VEGF Bridge Protein. J. Virol. 75, 1571–1575.

Sofroniew, M.V., 2020. Astrocyte reactivity: Subtypes, states, and functions in CNS innate immunity. Trends Immunol. 41, 758–770.

Stanley, G., Gokce, O., Malenka, R.C., Südhof, T.C., Quake, S.R., 2020. Continuous and discrete neuron types of the adult Murine striatum. Neuron 105, 688–699.e8.

Strebinger, D., Frangieh, C.J., Friedrich, M.J., Faure, G., Macrae, R.K., Zhang, F., 2023. Cell type-specific delivery by modular envelope design. Nat. Commun. 14, 5141.

Suzuki, T., Morimoto, N., Akaike, A., Osakada, F., 2020. Multiplex Neural Circuit Tracing With G-Deleted Rabies Viral Vectors. Front. Neural Circuits 13, 1–23.

Teufel, F., Almagro Armenteros, J.J., Johansen, A.R., Gíslason, M.H., Pihl, S.I., Tsirigos, K.D., Winther, O., Brunak, S., von Heijne, G., Nielsen, H., 2022. SignalP 6.0 predicts all five types of signal peptides using protein language models. Nat. Biotechnol. 40, 1023–1025.

Trinh, R., Gurbaxani, B., Morrison, S.L., Seyfzadeh, M., 2004. Optimization of codon pair use within the (GGGGS)3 linker sequence results in enhanced protein expression. Mol. Immunol. 40, 717–722.

Watakabe, A., Kato, S., Kobayashi, K., Takaji, M., Nakagami, Y., Sadakane, O., Ohtsuka, M., Hioki, H., Kaneko, T., Okuno, H., Kawashima, T., Bito, H., Kitamura, Y., Yamamori, T., 2012. Visualization of cortical projection neurons with retrograde TET-off lentiviral vector. PLoS One 7, e46157.

Yoshizawa, T., Ito, M., Doya, K., 2018. Reward-Predictive Neural Activities in Striatal Striosome Compartments. eNeuro 5. 10.1523/ENEURO.0367-17.2018

Yu, B., Shi, Q., Belk, J.A., Yost, K.E., Parker, K.R., Li, R., Liu, B.B., Huang, H., Lingwood, D., Greenleaf, W.J., Davis, M.M., Satpathy, A.T., Chang, H.Y., 2022. Engineered cell entry links receptor biology with single-cell genomics. Cell 185, 4904–4920.e22.

Zeng, H., 2022. What is a cell type and how to define it? Cell 185, 2739–2755.

